# Identifying a neurochemical signature of the paraventricular nucleus of the thalamus that predicts individual variability in cue-reward learning

**DOI:** 10.64898/2026.11.06.764855

**Authors:** Brianna A. Ramos, Sayan Chakrabarty, Caitlin N. Cain, Elizaveta Levina, Robert T. Kennedy, Shelly B. Flagel

## Abstract

To investigate neurochemical factors that may confer vulnerability to psychiatric disorders, we used an animal model that captures a transdiagnostically relevant neurobehavioral endophenotype. During a Pavlovian conditioned approach (PavCA) paradigm, rats learn the predictive relationship between a lever-cue and food reward. A subset of rats, sign-trackers (STs), also attribute incentive motivational value to the lever-cue. Relative to non-sign-trackers (NSTs), STs are more impulsive, have attentional deficits, and show greater reinstatement of drug-seeking behavior. The paraventricular nucleus of the thalamus (PVT) is a key neural node implicated in these individual differences. We combined in vivo microdialysis with liquid chromatography–tandem mass spectrometry (LC-MS/MS) to quantify 22 neurobiologically relevant small molecules in the PVT before and 24 hours after PavCA training. Before training, PVT glucose concentrations were higher in rats subsequently classified as STs than in NSTs and were correlated with the PavCA index used to phenotype the rats. Glucose concentrations decreased after training, and the magnitude of change differed between phenotypes. Multivariate regression identified baseline PVT neurochemical profiles associated with subsequent PavCA behavior. These findings identify a candidate PVT neurochemical signature that precedes behavioral phenotyping and may mark the propensity to sign-track, providing insight into potential mechanisms underlying vulnerability to psychiatric disorders characterized by maladaptive cue-motivated behavior.

## Introduction

Environmental stimuli, or cues, can guide adaptive behavior by directing individuals toward valuable resources (e.g., food and water) or away from danger. However, through Pavlovian learning and underlying neurobiological processes, reward-associated cues can acquire excessive control over behavior and promote maladaptive actions^1-4^. For example, exposure to a cue associated with drug use can trigger craving and relapse, due in part to altered dopamine signaling in the brain^5-7^. Similarly, a food-associated cue, such as the well-known golden arches of McDonald’s, can evoke food seeking and consumption even in the absence of physiological need. Although such cues are not inherently maladaptive, they can elicit excessive craving or desire in susceptible individuals and thereby contribute to maladaptive behavior^1,2,5,8^.

A Pavlovian conditioned approach (PavCA) paradigm can capture individual variation in cue-reward associative learning, enabling the study of neural processes relevant to psychiatric disorders, including addiction^2,9^. Following PavCA training, rats differ in whether their conditioned responses are primarily directed toward a reward-predictive cue or the site of reward delivery^3^. Sign-trackers (STs) approach and interact with the cue, whereas goal-trackers (GTs) preferentially approach the location of reward delivery. For both phenotypes, the cue acquires predictive value and elicits a conditioned response. For STs, however, the cue also acquires incentive motivational value, becoming attractive and desirable in its own right^3^.

Relative to GTs, STs are more susceptible to behavioral control by both food- and drug-paired cues^10-12^. STs and GTs represent opposite extremes of behavioral responding, whereas intermediate responders (INs) vacillate between these responses. In the present study, INs and GTs were collectively classified as non-sign-trackers (NSTs), encompassing rats that did not meet the predefined criterion for sign-tracking. This distinction allowed us to focus on neurochemical features associated with the propensity to sign-track, a phenotype characterized by the attribution of incentive salience to reward cues and increased susceptibility to their control over behavior^4^.

Individual differences in cue-motivated behavior have also been demonstrated in humans^13-16^. Greater sign-tracking tendencies have been associated with increased impulsivity, supporting the relevance of this phenotype to vulnerability to addiction and related psychiatric disorders^13,14^. Similarly, youth classified as STs exhibit greater externalizing behaviors and reduced inhibitory control compared with NST youth^15^. Functional neuroimaging further revealed greater reward-cue-elicited subcortical reactivity in ST youth, whereas NST youth more strongly engaged cortical control regions^15^. These findings suggest that sign-tracking may reflect a greater reliance on bottom-up subcortical motivational processes, whereas cue-motivated behavior in NST individuals may be more strongly regulated by top-down cortical control. This cross-species correspondence supports the translational relevance of the ST versus NST distinction and suggests that the balance between cortical control and subcortical cue reactivity may be a key neural mechanism underlying individual vulnerability to maladaptive cue-motivated behavior^15-18^.

Consistent with this proposed balance between cortical control and subcortical motivational processes, our laboratory has demonstrated in rats that top-down input from the prelimbic cortex (PrL) to the paraventricular nucleus of the thalamus (PVT) encodes the predictive value of reward cues and constrains bottom-up processes that promote incentive learning, including dopamine signaling in the nucleus accumbens (NAc)^19^. These findings led us to hypothesize that incentive learning associated with maladaptive motivated behavior emerges when bottom-up motivational processes supersede top-down control, with the PVT serving as a key integrative node. As a midline thalamic structure, the PVT is anatomically positioned to integrate information from arousal, limbic, cortical, subcortical, and motor circuits^20,21^. It has also been directly implicated in individual variation in Pavlovian conditioned responses^22^. The PVT receives dense glutamatergic input from the medial prefrontal cortex, predominantly the PrL ^23,24^, and sends glutamatergic projections to multiple cortical and subcortical regions, including the PrL, hypothalamus, hippocampus, amygdala, and NAc^24-29^. Through its position within hypothalamic-thalamic-striatal circuitry, the PVT contributes to cue-motivated behavior^21^. Indeed, Pavlovian reward cues elicit greater neuronal activation throughout this “motive circuit” in ST rats than in NST rats^30,31^.

Neuromodulators within the PVT contribute to homeostatic functions and motivated behavior^32-34^. For example, the PVT receives dense orexinergic input from the lateral hypothalamus^36,37^, and orexins excite PVT neurons^23,35^. Orexinergic signaling within the PVT has been implicated in arousal and motivated behavior^20^, including individual differences in cue-reward learning^40^. Orexin neurons have been shown to co-release glutamate in other target regions^38,39^, raising the possibility that orexinergic afferents also contribute to the extracellular profile of molecules detected in the PVT. However, the broader extracellular small-molecule neurochemical profile of the PVT remains poorly characterized. Defining this profile and its relationship to the propensity to sign-track could identify neurochemical features associated with individual differences in cue-motivated behavior.

The present study aimed to address this gap by quantifying a panel of extracellular neurotransmitters, metabolites, and energy-related molecules in the PVT and determining how this neurochemical profile relates to individual variation in cue-motivated behavior. In vivo microdialysis was combined with liquid chromatography–tandem mass spectrometry (LC-MS/MS) to simultaneously quantify 22 small-molecule analytes in the PVT before and after PavCA training. This within-subject design enabled us to distinguish baseline neurochemical features associated with the subsequent propensity to sign-track from neurochemical changes associated with PavCA training. We hypothesized that PVT neurochemical profiles would differ between rats subsequently classified as STs and NSTs and that baseline neurochemical measures would be associated with subsequent PavCA performance. Because maladaptive cue-motivated behavior is relevant to multiple psychiatric disorders, identifying PVT neurochemical features associated with the propensity to sign-track may reveal underlying biological mechanisms and potential therapeutic targets.

## Results and Discussion

To determine whether PVT neurochemistry was associated with the propensity to sign-track or with changes following PavCA training, we integrated our behavioral approach with longitudinal microdialysis sampling and analytical chemistry. Specifically, in vivo microdialysis was used to sample the extracellular environment of the PVT, while LC-MS/MS enabled simultaneous quantification of 22 neurotransmitters, metabolites, and energy-related molecules. Using a within-subject design (Figure 1), we characterized each rat’s PVT neurochemical profile before PavCA training (Pre-PavCA, Day 0) and again 24 h after the final training session (Post-PavCA, Day 6). Neurochemical concentrations could therefore be evaluated as a function of training stage and behavioral phenotype (ST versus NST). Importantly, because phenotype was determined only after training, associations between Pre-PavCA neurochemical concentrations and subsequent PavCA performance identified baseline features associated with the propensity to sign-track. In contrast, Pre-to Post-PavCA differences identified neurochemical changes associated with PavCA training. An additional group of control (Ctrl) animals received unpaired or randomized lever and food presentations, as previously described^30,31^. Histological verification confirmed that microdialysis probes were located predominantly within the medial and posterior PVT (Figure 1d; see also Supporting Figure 1).

**Figure 1.**
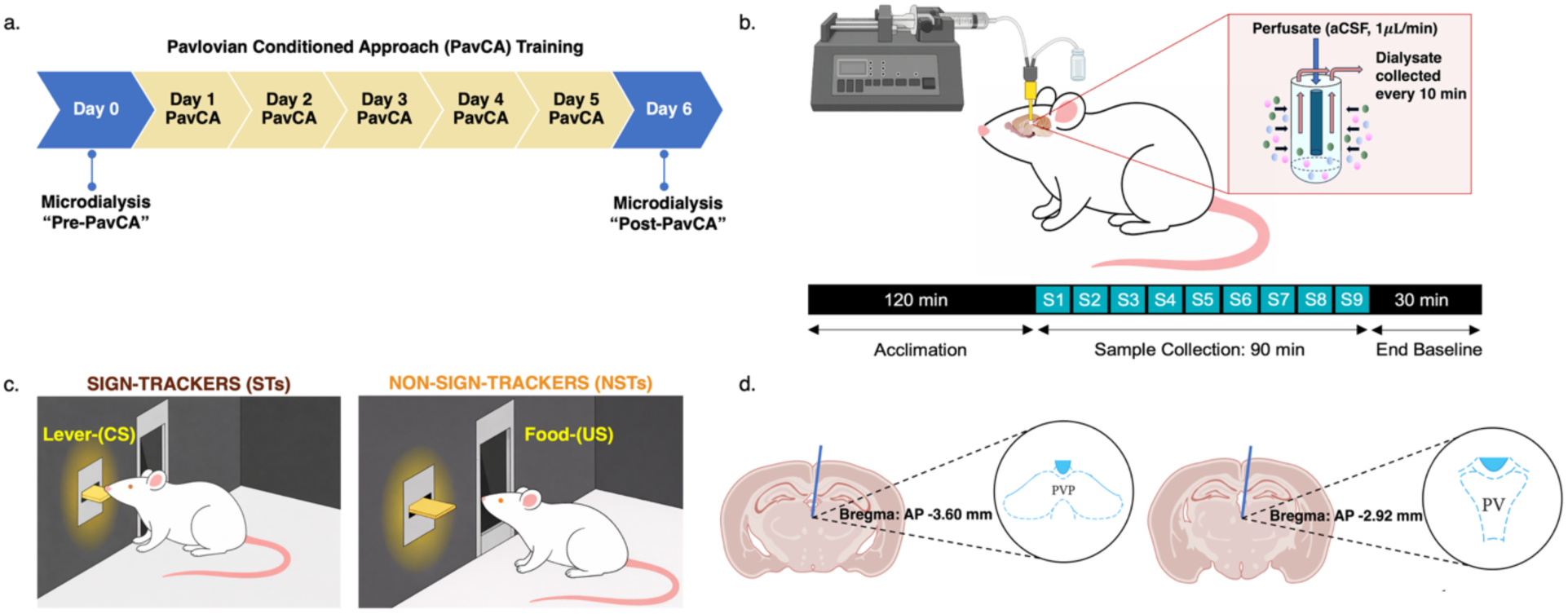
Experimental procedures. **(a)** Timeline of microdialysis collection: baseline sampling on Day 0 prior to Pavlovian conditioned approach training (Pre-PavCA), followed by five PavCA acquisition sessions, and concluding with post-training microdialysis collection on Day 6 (Post-PavCA). **(b)** Schematic of in vivo microdialysis sampling. Artificial cerebrospinal fluid (aCSF) was delivered via a syringe pump (1μL/min) into the inlet of the microdialysis probe. Small molecules passively diffused across the semipermeable membrane and were collected from the outlet. Dialysate was collected in 10-minute fractions over a 90-minute period. **(c)** Schematic of the PavCA training paradigm, as described in the text, with the behavior of sign-trackers (ST; left) and non-sign-trackers (NST; right) depicted. **(d)** Atlas-based representation of microdialysis probe placements in the paraventricular nucleus of the thalamus (PVT, referred to as PV in the atlas image). Majority of probe placements were located within a Bregma range of AP:−2.92 mm to AP:−3.60 mm, with the overall range from AP:−2.16 mm to AP:−4.08 mm, all within the medial through posterior PVT (pPVT, referred to as PVP in atlas image). Figures were created or adapted using BioRender. Sagittal brain image by Gill Brown (King’s College London). Brain atlas adapted from George Paxinos and Charles Watson 6^th^ edition 2007^60^. Sprague Dawley rat image adapted from iStock (Alina Umarova).

### PavCA training reveals individual differences in cue-directed behaviors

Male and female Sprague-Dawley rats underwent either five PavCA training sessions or five unpaired cue-reward sessions. PavCA-trained rats were categorized according to their PavCA index^41^ and the conditioned response that emerged across training. Rats receiving unpaired cue-reward sessions served as the control (Ctrl) group and were run alongside PavCA-trained animals. Because rats in the Ctrl group did not develop a conditioned response, they were not included in the behavioral analyses. STs exhibited behavior directed toward the lever-cue (Figure 2a-c, Table 1a-c), whereas NSTs exhibited behavior directed toward the location of reward delivery or showed little preference for either the lever-cue or reward location (Figure 2d-f, Table 1d-f). Relative to NSTs, STs exhibited greater lever-cue-directed behavior across training, including more contacts, a greater probability of contacting, and a shorter latency to contact the lever-cue. For each lever-directed measure, there were significant effects of Session and Phenotype and a significant Session x Phenotype interaction (all p<0.0001; Figure 2a-c, Table 1a-c).

**Figure 2:**
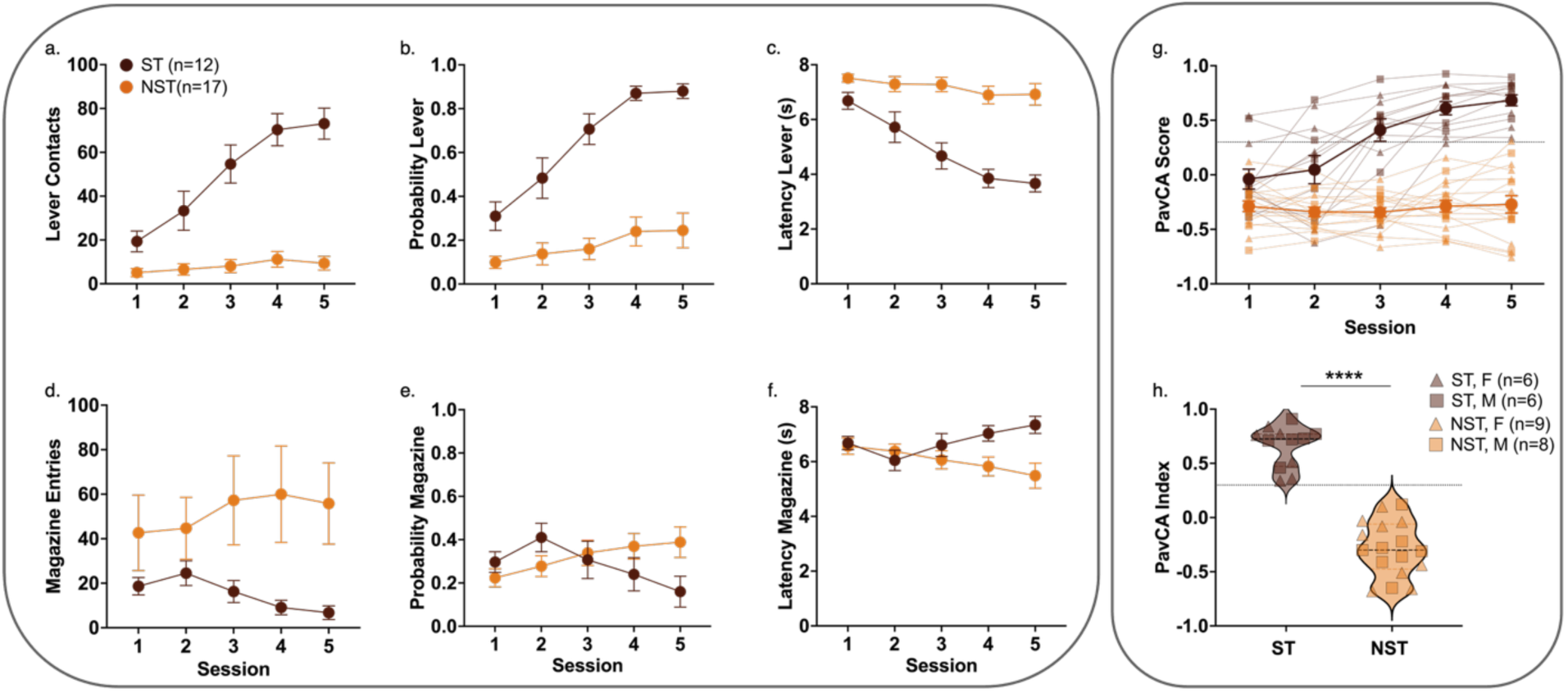
Behavioral characterization of sign-tracker (ST) and non-sign-tracker (NST) rats across 5 sessions of Pavlovian conditioned approach (PavCA) training. PavCA behavior is shown for (a–c) acquisition of lever-directed behaviors (sign-tracking) and (d-f) magazine-directed behavior (non-sign-tracking) during 5 PavCA sessions for sign-trackers (brown, ST, n=12) and non-sign-trackers (orange, NST, n=17). Mean ± SEM are illustrated for (a) number of lever-cue contacts, (b) probability of contacting the lever-cue, and (c) latency to contact the lever-cue, (d) number of magazine entries, (e) probability of entering the magazine, and (f) latency to enter the magazine. (g) PavCA score reflects a composite metric of lever-vs. magazine-directed behaviors for individual rats across 5 PavCA sessions. Data are shown for individual rats (lighter colors, smaller data points) with darker lines and larger data points representing the group mean ± SEM. (h) Violin plot showing the PavCA index (average of PavCA score across sessions 4 and 5) with the first quartile, median, and third quartile shown as dashed lines for male (M, squares) and female (F, triangles) sign-trackers (ST) and non-sign-trackers (NSTs); unpaired t-test (****p<0.0001). (g, h) Dotted black line at 0.3, indicates the phenotype threshold for STs and NSTs, with ≥0.3 representing STs and <0.3 representing NSTs.

**Table 1.**
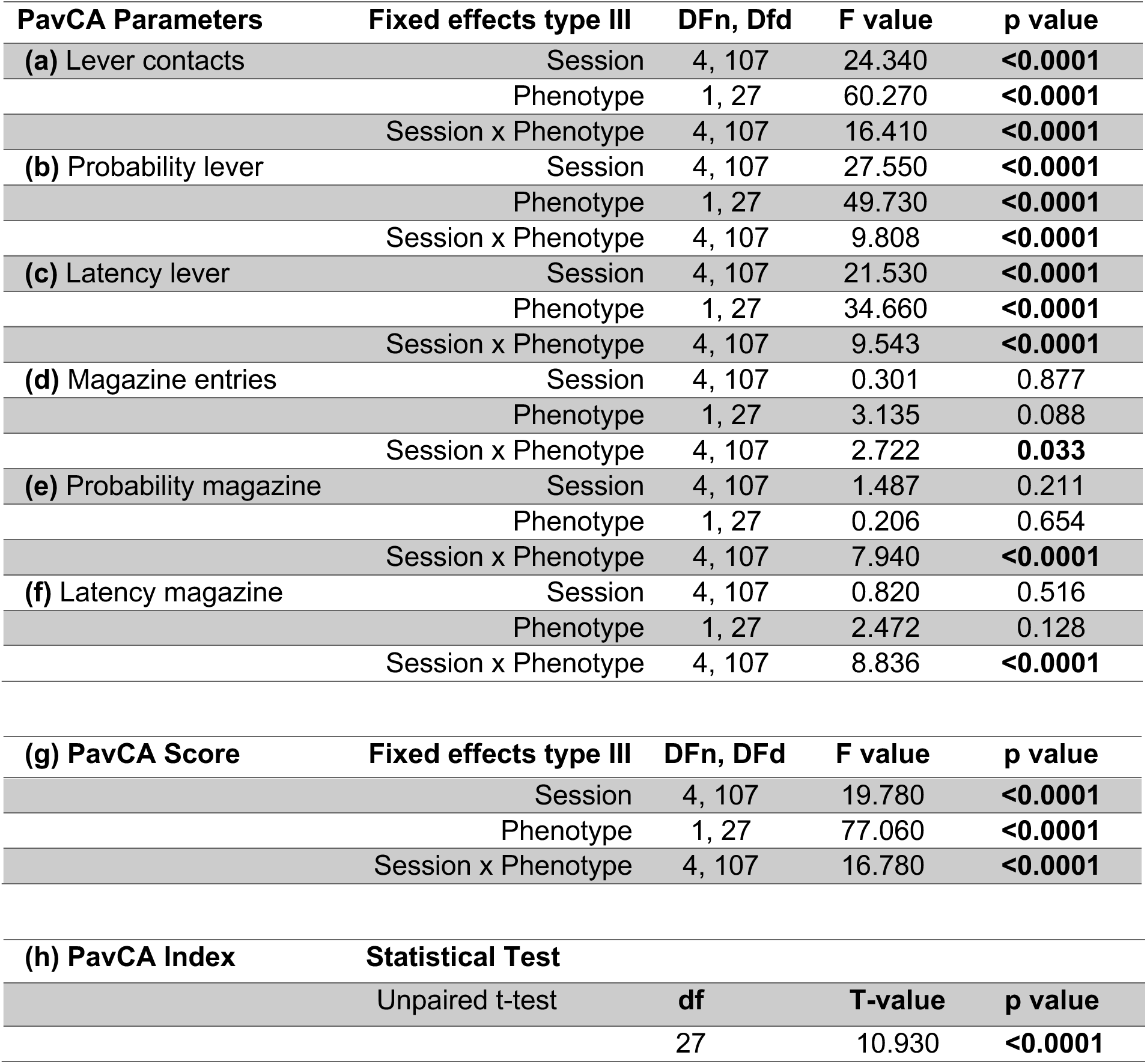
PavCA Behavior-Manuscript Figure 2. ***Statistical analyses of behavior during Pavlovian conditioned approach (PavCA) training.*** Mixed-effects models with Session, Phenotype (ST versus NST), and their interaction as fixed effects were used to analyze **(a)** lever-cue contacts, **(b)** probability of contacting the lever-cue, **(c)** latency to contact the lever-cue, **(d)** magazine entries, **(e)** probability of entering the magazine, **(f)** latency to enter the magazine, and **(g)** PavCA score. **(h)** An unpaired t-test compared PavCA indices between STs (n=12) and NSTs (n=17). Reported statistics include the applicable numerator and denominator degrees of freedom (DFn, DFd), F-values, t-values, and p-values. Panel labels correspond to Figure 2. Bold indicates p<0.05.

| PavCA Parameters | Fixed effects type III | DFn, Dfd | F value | p value |
|---|---|---|---|---|
| **(a)** Lever contacts | Session | 4, 107 | 24.340 | **<0.0001** |
|  | Phenotype | 1, 27 | 60.270 | **<0.0001** |
|  | Session x Phenotype | 4, 107 | 16.410 | **<0.0001** |
| **(b)** Probability lever | Session | 4, 107 | 27.550 | **<0.0001** |
|  | Phenotype | 1, 27 | 49.730 | **<0.0001** |
|  | Session x Phenotype | 4, 107 | 9.808 | **<0.0001** |
| **(c)** Latency lever | Session | 4, 107 | 21.530 | **<0.0001** |
|  | Phenotype | 1, 27 | 34.660 | **<0.0001** |
|  | Session x Phenotype | 4, 107 | 9.543 | **<0.0001** |
| **(d)** Magazine entries | Session | 4, 107 | 0.301 | 0.877 |
|  | Phenotype | 1, 27 | 3.135 | 0.088 |
|  | Session x Phenotype | 4, 107 | 2.722 | **0.033** |
| **(e)** Probability magazine | Session | 4, 107 | 1.487 | 0.211 |
|  | Phenotype | 1, 27 | 0.206 | 0.654 |
|  | Session x Phenotype | 4, 107 | 7.940 | **<0.0001** |
| **(f)** Latency magazine | Session | 4, 107 | 0.820 | 0.516 |
|  | Phenotype | 1, 27 | 2.472 | 0.128 |
|  | Session x Phenotype | 4, 107 | 8.836 | **<0.0001** |
| **(g) PavCA Score** | **Fixed effects type III** | **DFn, DFd** | **F value** | **p value** |
|  | Session | 4, 107 | 19.780 | **<0.0001** |
|  | Phenotype | 1, 27 | 77.060 | **<0.0001** |
|  | Session x Phenotype | 4, 107 | 16.780 | **<0.0001** |
| **(h) PavCA Index** | **Statistical Test** |  |  |  |
|  | Unpaired t-test | **df** | **T-value** | **p value** |
|  |  | 27 | 10.930 | **<0.0001** |

The trajectories of magazine-directed behavior also differed between STs and NSTs. By the end of training, NSTs made more magazine entries during lever-cue presentation (Session x Phenotype interaction, p=0.033; Figure 2d, Table 1d), had a greater probability of entering the food magazine (Session x Phenotype interaction, p<0.0001; Figure 2e, Table 1e), and exhibited a shorter latency to enter the magazine upon lever-cue presentation (Session x Phenotype interaction; p<0.0001, Figure 2f, Table 1f).

These behavioral differences were further reflected in the PavCA score^41^, a composite measure of response bias, probability, and latency that captures the relative tendency to direct behavior toward the lever-cue versus food magazine (Figure 2g, Table 1g). There were significant effects of Session (p<0.0001) and Phenotype (p<0.0001), as well as a Session x Phenotype interaction (p<0.0001; Table 1g). The PavCA index, calculated by averaging scores from the final two PavCA training sessions^41^, was used to classify rats as STs (n=12; index ≥0.3) or NSTs (n=17; index <0.3). Consistent with these classification criteria, PavCA indices differed significantly between STs and NSTs (p<0.0001; Figure 2h, Table 1h).

These results are indicative of both groups acquiring the predictive cue-reward association but differing in the incentive motivational value attributed to the lever-cue. All rats consumed all food rewards delivered, indicating that phenotypic differences were not attributable to differences in reward consumption. The resulting ST and NST phenotypes thus provided a behavioral framework for determining whether the propensity to sign-track was associated with distinct baseline or training-related neurochemical profiles in the PVT.

### PVT glucose concentrations are associated with the propensity to sign-track and change following PavCA training

Among the neurochemicals measured, glucose (Glc) showed the most consistent phenotype-associated differences and was reliably detected throughout the 90-min sampling period. We therefore examined Glc as an energy-related small molecule of particular relevance to the ST versus NST distinction. Figures 3a-b and Table 2a-b show extracellular PVT Glc concentrations across the 90-min sampling period before PavCA training, when the behavioral phenotype was not yet known, and after PavCA training, when rats had been classified as STs or NSTs. Glc concentrations were also examined for Ctrl animals before and after five unpaired cue-reward sessions (Figure 3c, Table 2c). Glc concentrations did not significantly change across the 90-min sampling period for either phenotype or the control group. Within each phenotype, however, there was a significant effect of Time Point (Pre-versus Post-PavCA), with lower concentrations after training in STs (p=0.0003; q=0.020; Figure 3a, Table 2a) and NSTs (p=0.028; q=0.144; Figure 3b, Table 2b). Glc concentrations also decreased in the Ctrl group following unpaired cue-reward sessions (p=0.030; q=0.338; Figure 3c, Table 2c). Although all groups showed a decrease in Glc concentration, only the decrease in STs remained significant after p-value correction (q-value). Corresponding analyses of additional small molecules are shown in Supporting Table 1.

**Figure 3.**
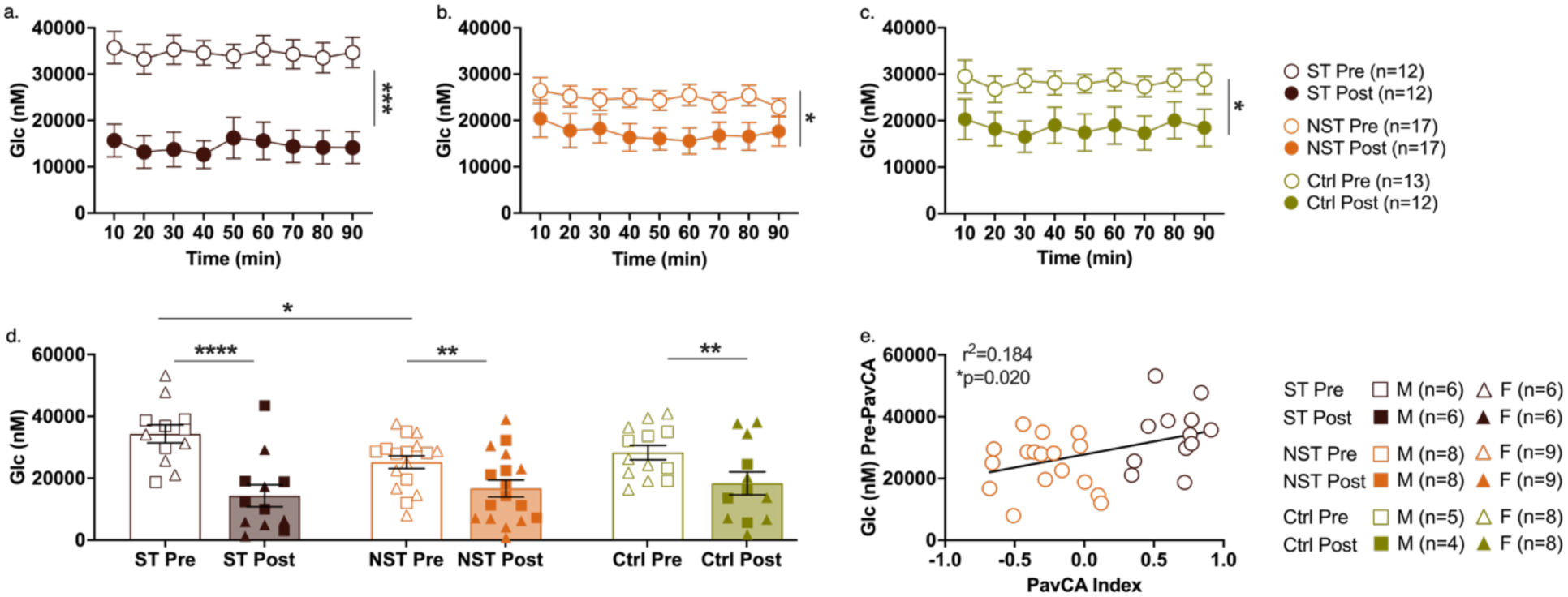
Glucose (Glc) concentration in the PVT before (Pre-) and after (Post-) Pavlovian Conditioned Approach (PavCA) training or unpaired cue-reward sessions for controls. (a-c) Mean ± SEM Glc concentrations (nM) measured by in vivo microdialysis across a 90-minute sampling period (nine 10-minute fractions) in (a) sign-tracker (ST) and (b) non-sign-tracker (NST) rats before and after PavCA training, or (c) control (Ctrl) animals which were presented with unpaired lever-cue and food reward. Mixed-effects analyses identified a significant effect of Time Point from Pre- to Post-PavCA for (a) ST (***p=0.0003; q=0.020), (b) NST (*p=0.028), and **(c)** Ctrl rats (*p=0.030). Asterisk indicate significance based on uncorrected p-value, except for STs, which remained significant after correction. **(d)** Mean ± SEM Glc concentrations averaged across nine sampling fractions shown for all three groups of animals. Individual data points indicate male (M) and female (F) rats. Mixed-effects analyses identified a significant main effect of Time Point (****p<0.0001; q=0.003) and a Time Point × Group interaction (*p=0.021). Fisher’s LSD multiple comparisons revealed a significant difference between Pre-and Post-Time Points for STs (****p<0.0001), NSTs (**p=0.003), and Ctrls (**p=0.002). STs and NSTs significantly differed in Glc concentration Pre-PavCA training (*p=0.024). **(e)** There was a significant correlation observed between Pre-PavCA Glc concentrations and PavCA index (*p=0.020). Corresponding analyses are shown in Table 2.

**Table 2.** Glucose Analyses - Manuscript Figure 3. ***Statistical analyses of PVT glucose concentrations before and after PavCA training or unpaired cue-reward sessions.*** (a–c) Glucose concentrations across the 90-min sampling period were analyzed separately within (a) sign-trackers (STs, n=12), (b) non-sign-trackers (NSTs, n=17), and (c) controls (Ctrls, n=13) using mixed-effects models with Sampling Period, Time Point (Pre versus Post), and their interaction as fixed effects. Pre and Post refer to measurements before and after PavCA training for STs and NSTs, or unpaired cue-reward sessions for Ctrls. (d) Mean glucose concentrations averaged across the 90-min sampling period were analyzed using a mixed-effects model with Time Point, Group (ST, NST, Ctrl), and their interaction as fixed effects. (e) Simple linear regression assessed the association between Pre-PavCA glucose concentration and subsequent PavCA index. Reported statistics include the applicable numerator and denominator degrees of freedom (DFn, DFd), F-values, p-values, Benjamini–Hochberg-adjusted q-values controlled across the 22 analytes (Supporting Table 2), and R². Unadjusted p-values <0.05 and q-values <0.05 are bolded.

| Groups analyzed | Fixed effects (type III) | DFn, DFd | F value | p value | q value |
|---|---|---|---|---|---|
| **(a)** ST Pre to Post | Sampling Period | 8, 173 | 1.568 | 0.138 | 0.420 |
|  | Time Point | 1, 22 | 19.010 | **0.0003** | **0.020** |
|  | Sampling Period x Time Point | 8, 173 | 0.675 | 0.713 | 0.898 |
| **(b)** NST Pre to Post | Sampling Period | 8, 239 | 1.500 | 0.158 | 0.521 |
|  | Time Point | 1, 32 | 5.276 | **0.028** | 0.144 |
|  | Sampling Period x Time Point | 8, 239 | 0.673 | 0.715 | 0.912 |
| **(c)** Ctrl Pre to Post | Sampling Period | 8, 169 | 1.389 | 0.204 | 0.627 |
|  | Time Point | 1, 23 | 5.370 | **0.030** | 0.338 |
|  | Sampling Period x Time Point | 8, 169 | 0.713 | 0.680 | 0.991 |
| **(d)** ST vs NST vs Ctrl | Time Point | 1, 38 | 57.180 | **<0.0001** | **0.003** |
|  | Group | 2, 39 | 0.531 | 0.592 | 0.759 |
|  | Time Point x Group | 2, 38 | 4.273 | **0.021** | 0.233 |
| **Groups analyzed** | **Simple linear regression** | **DFn, DFd** | **R<sup>2</sup> value** | **p value** | **q value** |
| **(e)** Pre-PavCA concentration vs PavCA index |  | 1, 27 | 0.184 | **0.020** | 0.339 |

When concentrations were averaged across the 90-min sampling period, mixed-effects analysis identified a significant main effect of Time Point (Pre-versus Post-PavCA; p<0.0001; q=0.003) and a significant Time Point x Group interaction (p=0.021; q=0.233; Figure 3d, Table 2d), suggesting that the magnitude of Pre to Post change differed across groups. Post hoc comparisons using uncorrected Fisher’s least significant difference (LSD) tests identified significant Pre to Post decreases in the average Glc concentration in each group (ST: p<0.0001; NST: p=0.003; Ctrl: p=0.002; Figure 3d). These comparisons also revealed higher Pre-PavCA Glc concentrations in rats subsequently classified as STs than in NSTs (p=0.024; Figure 3d). Corresponding analyses of mean concentrations by Time Point and Group for additional small molecules are shown in Supporting Table 2.

These findings indicate that differences in PVT Glc were present before rats underwent cue-reward associative learning. Moreover, Pre-PavCA Glc concentration was associated with the subsequent PavCA index (R^2^=0.184, p=0.020; q=0.339; Figure 3e, Table 2e), accounting for approximately 18% of variance in PavCA index, although this association did not remain significant after p-value correction. No other analytes showed a statistically significant association at the unadjusted threshold (Supporting Table 3).

**Table 3.** Analyte Analyses - Manuscript Figure 4. ***Statistical analyses of mean neurochemical concentrations in the PVT by Time Point and Phenotype.*** Neurochemical concentrations (nM) averaged across the 90-min sampling period were analyzed using mixed-effects models with Time Point (Pre-versus Post-PavCA), Phenotype (ST (n=12) versus NST (n=17)), and their interaction as fixed effects. Panel labels correspond to Figure 4a–n. Reported statistics include numerator and denominator degrees of freedom (DFn, DFd), F-values, p-values, and Benjamini–Hochberg-adjusted q-values controlled across the 22 analytes (Supporting Table 4). Unadjusted p-values <0.05 and q-values <0.05 are bolded. For certain analytes, including Asp, Ser, NE, NM, DA and 3MT, some values were missing due to weak signal or concentration falling below the detection threshold.

| Analyte | Fixed effects type III | DFn, DFd | F value | p value | q value |
|---|---|---|---|---|---|
| **(a)** Asp | Time Point | 1, 17 | 0.598 | 0.450 | 0.695 |
|  | Phenotype | 1, 26 | 1.119 | 0.300 | 0.641 |
|  | Time Point × Phenotype | 1, 17 | 1.723 | 0.207 | 0.641 |
| **(b)** Glu | Time Point | 1, 27 | 0.423 | 0.521 | 0.716 |
|  | Phenotype | 1, 27 | 0.002 | 0.963 | 0.973 |
|  | Time Point × Phenotype | 1, 27 | 0.553 | 0.464 | 0.695 |
| **(c)** Tau | Time Point | 1, 27 | 0.120 | 0.732 | 0.818 |
|  | Phenotype | 1, 27 | 1.941 | 0.175 | 0.578 |
|  | Time Point × Phenotype | 1, 27 | 0.070 | 0.793 | 0.872 |
| **(d)** Ser | Time Point | 1, 20 | 3.949 | 0.061 | 0.365 |
|  | Phenotype | 1, 27 | 2.066 | 0.162 | 0.563 |
|  | Time Point × Phenotype | 1, 20 | 0.223 | 0.642 | 0.768 |
| **(e)** Gly | Time Point | 1, 27 | 1.303 | 0.264 | 0.641 |
|  | Phenotype | 1, 27 | 1.319 | 0.261 | 0.641 |
|  | Time Point × Phenotype | 1, 27 | 0.020 | 0.888 | 0.945 |
| **(f)** GABA | Time Point | 1, 27 | 8.083 | **0.008** | 0.136 |
|  | Phenotype | 1, 27 | 0.208 | 0.652 | 0.768 |
|  | Time Point × Phenotype | 1, 27 | 0.757 | 0.392 | 0.673 |
| **(g)** Ado | Time Point | 1, 27 | 0.005 | 0.946 | 0.973 |
|  | Phenotype | 1, 27 | 3.659 | 0.066 | 0.365 |
|  | Time Point × Phenotype | 1, 27 | 0.288 | 0.596 | 0.755 |
| **(h)** NE | Time Point | 1, 50 | 0.905 | 0.346 | 0.641 |
|  | Phenotype | 1, 50 | 0.931 | 0.339 | 0.641 |
|  | Time Point × Phenotype | 1, 50 | 0.676 | 0.415 | 0.673 |
| **(i)** NM | Time Point | 1, 12 | 3.436 | 0.089 | 0.389 |
|  | Phenotype | 1, 20 | 8.024 | **0.010** | 0.136 |
|  | Time Point × Phenotype | 1, 12 | 2.307 | 0.155 | 0.563 |
| **(j)** DA | Time Point | 1, 25 | 1.165 | 0.291 | 0.641 |
|  | Phenotype | 1, 27 | 1.029 | 0.319 | 0.641 |
|  | Time Point × Phenotype | 1, 25 | 0.363 | 0.552 | 0.724 |
| **(k)** 3MT | Time Point | 1, 6 | 0.515 | 0.500 | 0.702 |
|  | Phenotype | 1, 16 | 6.229 | **0.024** | 0.197 |
|  | Time Point × Phenotype | 1, 6 | 0.141 | 0.720 | 0.818 |
| **(l)** HVA | Time Point | 1, 27 | 0.905 | 0.350 | 0.641 |
|  | Phenotype | 1, 27 | 1.115 | 0.300 | 0.641 |
|  | Time Point × Phenotype | 1, 27 | 0.241 | 0.628 | 0.767 |
| **(m)** Glc | Time Point | 1, 27 | 40.370 | **<0.0001** | **0.003** |
|  | Phenotype | 1, 27 | 1.095 | 0.305 | 0.641 |
|  | Time Point × Phenotype | 1, 27 | 6.589 | **0.016** | 0.152 |
| **(n)** Gln | Time Point | 1, 27 | 9.031 | **0.006** | 0.125 |
|  | Time Point × Phenotype | 1, 27 | 2.895 | 0.100 | 0.414 |

The difference in Glc concentration before PavCA training is particularly noteworthy because these measurements were obtained before rats experienced the cue-reward association. Thus, differences in PVT Glc could not be attributed to PavCA training. Rather, the association between Pre-PavCA Glc concentration and the subsequent PavCA index suggests that the basal extracellular metabolic environment of the PVT may represent a pre-existing neurochemical feature associated with the propensity to sign-track. This relationship warrants further investigation, both to determine whether baseline Glc availability causally contributes to the propensity to sign-track and to assess whether Glc is part of a broader metabolic signature associated with this propensity. Examining Glc together with related metabolites could clarify whether a coordinated PVT metabolic profile rather than a single molecule is associated with individual differences in cue-motivated behavior.

Extracellular brain Glc concentrations are related to the energetic demands of neural activity and reflect the balance among local Glc delivery, uptake, and utilization^42^. The differences observed here may therefore reflect variation in the functional or metabolic state of the PVT. This possibility is particularly relevant because the PVT is an integrative node within the circuitry underlying motivated behavior and sends dense projections to the NAc^21,25,26,28^. Previous work identified a population of glucose-responsive, glutamatergic, GLUT2-expressing PVT neurons that project to the NAc and regulate sucrose-seeking behavior^43^. Specifically, reduced Glc availability increased the activity of these “hypoglycemia-activated” neurons, and stimulation of their axons in the NAc elicited excitatory postsynaptic currents in NAc medium spiny neurons^43^. Together, these findings identify a potential pathway through which metabolic sensing in the PVT may influence the motive circuit. However, the present study cannot distinguish the relative contributions of neuronal Glc utilization, astrocytic metabolism, vascular delivery, or other processes that regulate local Glc availability^44^, nor can it identify the cells or pathways responsible for the observed Glc differences. Nonetheless, this prior work supports a biologically plausible relationship between PVT Glc availability and motivated behavior.

The significant decrease in Glc concentration from Pre-to Post-PavCA indicates that the baseline extracellular metabolic environment of the PVT changed following training. Although Glc concentrations also decreased in Ctrl animals following unpaired cue-reward sessions, the Time Point x Group interaction suggests that the magnitude of this change varied across groups. Furthermore, the association between Pre-PavCA Glc concentrations and subsequent PavCA index links baseline PVT Glc to the propensity to sign-track. Extracellular brain Glc has previously been shown to change in response to cognitive and behavioral demands^45^, although such changes have not been well characterized in the PVT^46^. Post-PavCA samples were collected 24 hours after the final training session; therefore, these differences are unlikely to reflect an acute response to the PavCA task and may instead represent persistent neurochemical changes. Given that both phenotypes learned the cue-reward association but differed in the motivational value attributed to the cue, these changes may reflect neural or metabolic processes associated with distinct cue-reward learning styles. The decrease observed in Ctrl animals suggests that processes other than associative learning may also contribute.

### PavCA training and behavioral phenotypes are associated with selective differences in PVT neurochemistry

Given the effects observed for Glc, we next examined whether PavCA training and behavioral phenotypes were associated with broader neurochemical differences within the PVT (Figure 4, Table 3). Neurochemical concentrations were averaged across the 90-min microdialysis sampling period and examined as a function of Time Point (Pre-versus Post-PavCA), Phenotype (ST versus NST), and the Time Point x Phenotype interaction.

**Figure 4.**
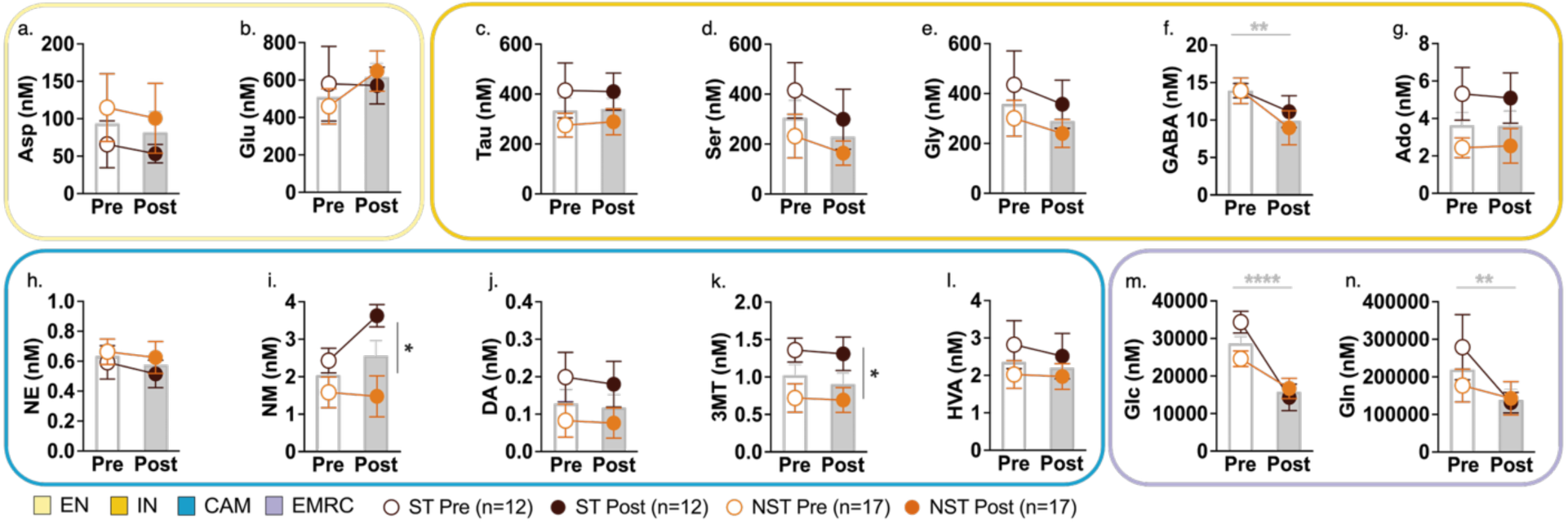
Neurochemical analytes measured within the PVT using in vivo microdialysis sampling Pre- and Post-PavCA training. **(a-n)** Mean ± SEM concentration (nM) of analytes averaged across the 90-min sampling period are shown before (Pre, white bars) and after (Post, gray bars) PavCA training for the entire population (n=29) and by phenotype (line, sign-tracker (ST), n=12 (brown); non-sign-tracker (NST), n=17 (orange)). Mixed-effects analyses revealed a significant effect of Time Point (Pre-vs. Post-; gray significant line and asterisks) for **(f)** *γ*-aminobutyric acid (GABA; **p=0.008), **(m)** glucose (Glc; ****p<0.0001, q=0.003; Time x Phenotype interaction *p=0.016) and **(n)** glutamine (Gln; **p=0.006). A significant effect of Phenotype (black asterisk, side) was observed for **(i)** normetanephrine (NM; *p=0.010) and **(k)** 3-methoxytyramine (3MT; *p=0.024). Asterisks indicate significance based on uncorrected p-values, except for Glc, which remained significant after correction. No significant effects were observed for **(a)** aspartate (Asp), **(b)** glutamate (Glu), or **(c)** taurine (Tau), **(d)** serine (Ser), **(e)** glycine (Gly), **(g)** adenosine (Ado), **(h)** norepinephrine (NE), **(j)** dopamine (DA), or **(l)** homovanillic acid (HVA). Neurochemical groups displayed are categorized and outlined as follows with corresponding colors: Excitatory Neurotransmitters (EN, light yellow), Inhibitory Neurotransmitters and Neuromodulators (IN, dark yellow), Catecholamines and Metabolites (CAM, cerulean blue), Energy Metabolism-Related Compounds (EMRC, light purple). Corresponding analyses are shown in Table 3.

Mixed-effects analyses revealed no significant effects of Time Point, Phenotype, or their interaction for excitatory neurotransmitters aspartate (Asp; Figure 4a, Table 3a) and glutamate (Glu; Figure 4b, Table 3b), or for taurine (Tau; Figure 4c, Table 3c), serine (Ser; Figure 4d, Table 3d), glycine (Gly; Figure 4e, Table 3e), or adenosine (Ado; Figure 4g, Table 3g). In contrast, there was a significant effect of Time Point for γ-aminobutyric acid (GABA; p=0.008; q=0.136; Figure 4f, Table 3f), which decreased Pre-to Post-PavCA.

Catecholamines and their metabolites are shown in Figure 4h-l and Table 3h-l. There was not a significant effect of Time Point, Phenotype, or their interaction for norepinephrine (NE; Figure 4h, Table 3h). However, STs tended to show higher concentrations than NSTs for the NE metabolite normetanephrine (NM; p=0.010; q=0.136; Figure 4i, Table 3i). Similarly, there were no significant effects for dopamine (DA; Figure 4j, Table 3j), but concentrations of the DA metabolite 3-methoxytyramine (3MT) tended to be greater in STs than in NSTs (p=0.0239; q=0.197; Figure 4k, Table 3k). Homovanillic acid (HVA) did not differ significantly between phenotypes (Figure 4l, Table 3l).

Lastly, concentrations of the metabolism-related compounds glucose (Glc; p<0.0001; q=0.003; Figure 4m, Table 3m) and glutamine (Gln; p=0.006; q=0.125; Figure 4n, Table 3n) decreased from Pre- to Post-PavCA, with Glc remaining significant after applying p-value correction. Consistent with the findings above, there was a significant Time Point x Phenotype interaction for Glc (p=0.016), suggesting phenotype-specific differences in the magnitude of the Pre to Post change. Corresponding analyses of additional analytes are included in Supporting Table 4. DOPEG also significantly changed from Pre- to Post-PavCA (p<0.0001; q=0.003), with a significant Time Point x Phenotype interaction (p=0.031); and histamine showed a similar trend (Time Point, p=0.015; q=0.152; Time Point x Phenotype, p=0.058, q=0.365).

Supporting Figure 2 shows corresponding data from the control animals, which were analyzed separately because they were not classified as STs or NSTs. In the mixed-effects analyses across the 90-min sampling period, nominally significant effects of Time Point were observed for GABA (p=0.030; q=0.338), NM (p=0.030; q=0.338), Glc (p=0.030; q=0.338), and Gln (p=0.011; q=0.338), but none remained significant after p-value correction (Supporting Table 1c; Table 2c for Glc). These results suggest that changes in neurochemical concentrations were not restricted to paired cue-reward training. However, overall Pre to Post changes in controls address a different question from the phenotype-associated effects, including a trend for higher NM and 3MT concentrations in STs than in NSTs and differences between phenotypes in the magnitude of the Pre to Post Glc change. Thus, the control findings do not negate these exploratory phenotype-associated differences, although they limit attribution of the overall Pre to Post changes specifically to associative learning.

The PVT is a heterogeneous brain region that receives neurochemically diverse inputs from cortical and subcortical regions^21^. Specifically, the PVT receives GABAergic input from subcortical regions, including the lateral hypothalamus (LH)^47^, a region implicated in cue-motivated behavior. Thus, the observed decrease in extracellular GABA from Pre-to Post-PavCA may reflect altered inhibitory regulation of the PVT following PavCA training. Given the involvement of LH-PVT circuitry in motivated behavior^47,48^, changes in GABAergic signaling from this or other afferent populations could contribute to the observed effect^49^. The concurrent decrease in Gln may reflect altered amino acid metabolism or neurotransmitter cycling following PavCA training^50,51^. The changes in GABA and Gln observed in control animals suggest that these Time Point effects may reflect processes shared by paired and unpaired cue-reward exposure, rather than processes specific to associative learning. The current study cannot identify the cellular or circuit-level processes underlying the observed extracellular changes in GABA and Gln. Future studies employing chemogenetic or optogenetic approaches to manipulate specific inputs to the PVT could help determine their causal contributions to training-associated neurochemical changes.

In contrast to the overall Time Point effects, the effects of Phenotype for NM and 3MT identified differences in catecholamine metabolite concentrations between STs and NSTs across the two sampling stages. These phenotype-associated differences remain informative even when an analyte, such as NM, also changes over time in control animals. Although extracellular NE and DA concentrations did not significantly differ between STs and NSTs, measurements of their metabolites provide information complementary to that obtained from the parent neurotransmitters. Differences in extracellular metabolite concentrations may reflect variation in neurotransmitter release, uptake, metabolism, clearance, or some combination of these processes^52,53^. Accordingly, NM and 3MT concentrations should not be interpreted as direct measures of catecholamine turnover but rather as phenotype-associated features of catecholamine-related neurochemistry within the PVT.

The PVT receives catecholaminergic input from several brain regions that could contribute to these phenotype-related differences. The locus coeruleus (LC), a major source of NE in the brain, projects to the PVT^54^, and LC terminals in the PVT can also release dopamine^55^. In contrast, dopaminergic fibers in the PVT arise primarily from dopamine cell groups in the hypothalamus and periaqueductal gray^56^. Additionally, PVT neurons are known to express dopamine D2 receptors, which is consistent with dopamine input^57^. These afferent systems represent potential circuit-level sources of the catecholamines whose metabolites differed between phenotypes. Additional studies are needed to determine whether these differences reflect altered activity within these pathways or local differences in catecholamine metabolism and clearance.

Consistent with the present findings, a previous microdialysis LC-MS/MS study from our laboratory identified phenotype-associated differences in nucleus accumbens catecholamine concentrations in selectively bred high-responder (bHR) and low-responder (bLR) rats, with bHR rats exhibiting higher basal DA and NE concentrations^52,58^. Although the animal model and brain region differ from those examined here, these findings provide convergent evidence that neurobehavioral phenotypes relevant to addiction^2,59^ are accompanied by differences in extracellular catecholamine and metabolite profiles.

### Multivariate PVT neurochemical profiles are associated with the PavCA index

Given the neurochemical differences observed between STs and NSTs, we next examined whether multivariate PVT neurochemical profiles were associated with individual variation in the PavCA index. This analysis addressed whether the combined neurochemical profile captured variation across the behavioral spectrum, beyond differences in individual analytes. Multiple linear regression models were generated separately using Pre- or Post-PavCA neurochemical concentrations, with the PavCA index treated as a continuous outcome. Bayesian information criterion (BIC)-based stepwise selection was used to identify reduced sets of analytes associated with the PavCA index. Model-derived PavCA indices were also used to classify rats as STs or NSTs using an index threshold of 0.3, based on the criteria described above. Model performance was evaluated using 1,000 stratified train-test splits to avoid optimistic bias from overfitting.

The Pre-PavCA BIC-selected model retained 12 analytes spanning several neurochemical classes: NM, Glc, Ser, 3MT, Ch, NE, Tau, HVA, DOPAC, VMA, DA, and Gln (Figure 5a-c). In the full dataset, the model accounted for 82.7% of the variation in PavCA index (R^2^=0.827), with a root mean squared error (RMSE) of 0.214 and a normalized root mean squared error (NRMSE) of 13.5%. Classification based on the model-derived PavCA index was 93.1% accurate, with two STs misclassified as NSTs. Both misclassified rats had observed PavCA indices near the classification threshold.

**Figure 5.**
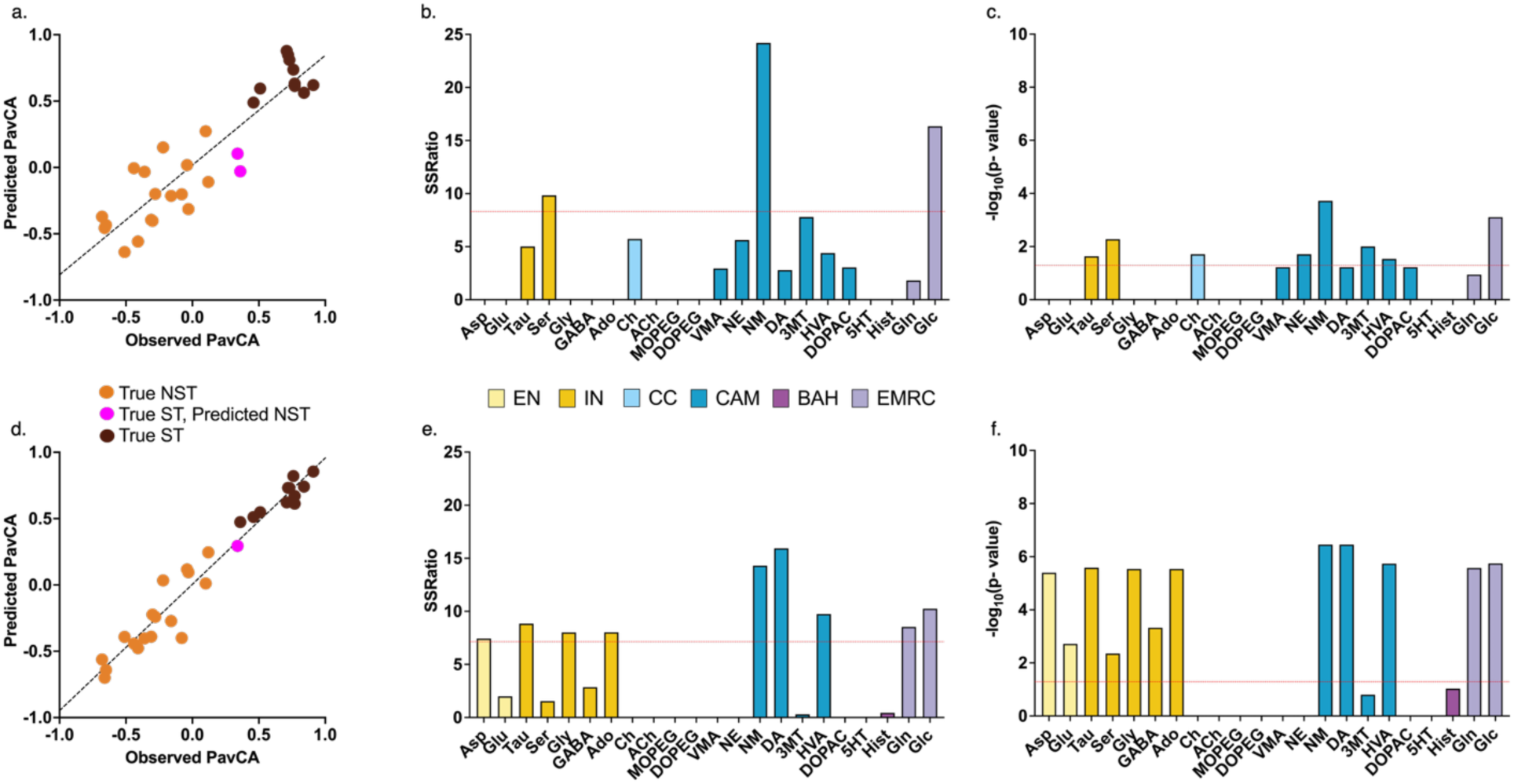
The ‘neurosignature’ of the PVT can be used as a predictor for PavCA index and behavioral phenotype. **(a–c)** Pre-PavCA and **(d–f)** Post-PavCA PVT neurochemical profiles were used to generate multiple linear regression models of PavCA index following Bayesian information criterion (BIC)-based stepwise selection. **(a, d)** Model-predicted versus observed PavCA index for the **(a)** Pre- and **(d)** Post-PavCA models. Model-predicted indices were used to classify rats as non-sign-trackers (NSTs; <0.3) or sign-trackers (STs; ≥0.3), with misclassified animals shown in pink. **(a)** Pre-PavCA model performance: R²=0.827, RMSE=0.214, NRMSE=0.135, and accuracy=93.1%; cross-validated performance: CV-R²=0.503, CV-RMSE=0.370, CV-NRMSE=0.309, and CV-accuracy=86.3%. **(d)** Post-PavCA model performance: R²=0.952, RMSE=0.113, NRMSE=0.071, and accuracy=96.6%; cross-validated performance: CV-R²=0.535, CV-RMSE=0.357, CV-NRMSE=0.295, and CV-accuracy=88.0%. **(b, e)** Predictor contributions are shown as sum of squares ratios (SSRatios), with red dashed lines indicating the mean SSRatio within each model. **(c, f)** Predictor significance is shown as −log10(*p*-values) from *t*-tests of regression coefficients; higher values indicate greater statistical significance. Red dashed lines indicate −log10(0.05), with values above the line corresponding to *p*<0.05. Bars are color-coded by neurochemical class: Excitatory Neurotransmitters (EN, light yellow), Inhibitory Neurotransmitters and Neuromodulators (IN, dark yellow), Cholinergic Compounds (CC, light blue), Catecholamines and Metabolites (CAM, blue), Biogenic amine– Histamine (BAH, dark purple), and Energy Metabolism-Related Compounds (EMRC, light purple). Abbreviations: dopamine (DA), norepinephrine (NE), serotonin (5-HT), histamine (Hist), glutamate (Glu), γ-aminobutyric acid (GABA), acetylcholine (ACh), choline (Ch), adenosine (Ado), 3,4-dihydroxyphenylacetic acid (DOPAC), 3-methoxytyramine (3MT), homovanillic acid (HVA), normetanephrine (NM), taurine (Tau), serine (Ser), aspartate (Asp), glycine (Gly), glutamine (Gln), vanillylmandelic acid (VMA), 3,4-dihydroxyphenylethylene glycol (DOPEG), 3-methoxy-4-hydroxyphenylethylene glycol (MOPEG), and glucose (Glc).

The Pre-PavCA model retained substantial information about subsequent PavCA behavior under cross-validation. Across 1,000 stratified train-test splits, mean cross-validated R^2^ was 0.503, RMSE was 0.370, NRMSE was 30.87%, and phenotype classification accuracy was 86.3%. These results suggest that baseline PVT neurochemistry captured individual differences relevant to the subsequent propensity to sign-track, before rats experienced paired cue-reward training.

To examine the contributions of individual analytes, predictor importance was assessed using the sum of squares ratio (SSRatio) and Benjamini-Hochberg adjusted p-values for individual regression coefficients (Figure 5b, Figure 5c). The analytes that were strongest contributors to the Pre-PavCA model were: Tau, Ser, Ch, HVA, NM, NE, 3MT and Glc. These analytes span amino acids, catecholamines and their metabolites, and energy metabolism-related compounds. This multivariate pattern suggests that subsequent PavCA behavior was associated with a distributed PVT neurochemical profile rather than with a single analyte. Notably, both energy metabolism-related compounds measured, Glc and Gln, were retained in the Pre-PavCA model, although Gln was not among the strongest contributors (Figure 5b-c). This finding is consistent with the possibility that Glc is associated with the propensity to sign-track as part of a broader metabolic profile rather than as an isolated neurochemical marker.

We next examined the Post-PavCA neurochemical profile. The BIC-selected model retained 14 analytes: DA, NM, Glc, HVA, Tau, Gln, Ado, Gly, Asp, GABA, Glu, Ser, Hist, and 3MT (Figure 5d-f). In the full dataset, the model accounted for 95.2% of the variation in the PavCA index (R^2^=0.952), with an RMSE of 0.113, NRMSE of 7.12%, and a phenotype classification accuracy of 96.6%. Under cross-validation, mean R^2^ was 0.535, RMSE was 0.357, NRMSE was 29.49%, and phenotype-classification accuracy was 88.0%. Thus, the Post-PavCA neurochemical profile also captured individual variation in PavCA behavior, with cross-validated performance comparable to that of the Pre-PavCA model.

Predictor-importance analyses indicated that DA, NM, Glc, HVA, Tau, Gln, Ado, Gly, Asp, GABA, Glu, and Ser contributed strongly to the Post-PavCA model (Figures 5e and 5f). NM, Glc, HVA, Tau, and Ser were among the strongest contributors to both the Pre- and Post-PavCA models, suggesting that these analytes may represent relatively consistent components of the PVT neurochemical profiles associated with PavCA behavior. The Post-PavCA model also included several excitatory and inhibitory neurochemicals, including Asp, Glu, Gly, GABA, and Ado. These differences in model composition may reflect training-associated changes in the relationships between PVT neurochemistry and PavCA behavior. Glc and Gln were again retained in the Post-PavCA model, with both contributing strongly, consistent with the possibility that energy-related molecules jointly reflect PVT metabolic states associated with individual differences in cue-reward learning. The reduced performance under cross-validation and potential instability of variable selection in this small sample warrant caution in interpreting individual predictor rankings and differences in model composition. Independent validation is needed to establish generalizability.

Together, these exploratory models identified multivariate PVT neurochemical profiles associated with individual differences in PavCA behavior. Importantly, the Pre-PavCA profile was associated with subsequent PavCA index, indicating that behaviorally relevant neurochemical variation was present before paired cue-reward training and behavioral classification. The Post-PavCA profile further indicates that individual differences in PavCA behavior remain associated with PVT neurochemistry after training. These associations address a different question from the overall Pre to Post changes observed in control animals, as they link variation in PVT neurochemistry to the propensity for and expression of sign-tracking among PavCA-trained rats. These findings support the existence of a candidate PVT “neurosignature” associated with individual differences in susceptibility to cue-motivated behavior.

## Conclusions

The present study characterized the extracellular neurochemical profile of the PVT in outbred male and female rats before and after PavCA training. Using longitudinal in vivo microdialysis coupled with LC-MS/MS, we identified baseline neurochemical features related to individual differences in cue-reward learning. Most notably, Pre-PavCA Glc concentrations were higher in rats subsequently classified as STs than in NSTs, indicating that this difference was present before behavioral phenotypes emerged. The magnitude of the Pre-to Post-PavCA decrease in Glc also differed between phenotypes, suggesting a persistent, phenotype-dependent shift in the extracellular metabolic environment of the PVT following training. Higher concentrations of the catecholamine metabolites NM and 3MT in STs further indicate that phenotype-associated differences extend beyond Glc. These phenotype-associated findings met the unadjusted significance threshold but remain exploratory because they did not survive p-value correction. Decreases in GABA and Gln following both paired and unpaired cue-reward sessions also revealed broader changes in PVT neurochemistry that were not specific to associative learning.

Exploratory multivariate modeling further showed that Pre-and Post-PavCA neurochemical profiles contained information associated with individual variation in the PavCA index, supporting the value of examining neurochemical systems collectively rather than focusing on isolated analytes. These findings support candidate PVT “neurosignatures” associated with the propensity for and expression of sign-tracking. Validation in larger, independent, sex-balanced cohorts will be necessary to establish their predictive utility and determine whether these relationships differ by sex, a question the present study was not powered to address. Collectively, these findings identify a multivariate PVT neurochemical profile associated with the subsequent propensity to sign-track and the expression of individual differences in cue-reward learning. This work provides a foundation for investigating neurochemical mechanisms that may contribute to vulnerability to maladaptive cue-driven behavior relevant to psychiatric disorders.

## Methods

All procedures conformed to *The Guide for the Care and Use of Laboratory Animals* and were approved by the University of Michigan Institutional Animal Care and Use Committee. Figure 1 illustrates the experimental timeline, with details described below.

### Subjects

Male (n=39) and female (n=39) Sprague-Dawley rats were purchased from Charles River (Stone Ridge, NY; barriers R08, R09, R04, K90, K98) and arrived at our facilities at approximately 60 days old, weighing 250-300g on arrival. Rats had free access to food and water and were maintained on a 12:12 light/dark cycle (lights on at 7:00 am) for the duration of the study. Rats were initially housed in same-sex pairs and then individually housed following cannulation surgeries. All experimentation occurred during the light cycle, between the hours of 8:00 am and 12:30 pm. Data were collected across six cohorts of rats. Due to laboratory relocation, three of the six cohorts were housed in a different vivarium facility, but all rats were maintained under the same housing and experimental conditions. Attrition occurred due to surgical complications, inaccurate probe placement, and technical issues, resulting in a total sample size of 42 rats (n=19 males, n=23 females; n=29 PavCA-trained rats, n=13 Ctrl rats).

### Stereotaxic surgeries for microdialysis probe placement

Stereotaxic surgeries for microdialysis guide cannulae occurred approximately 1 week after rats arrived in our facility. Rats were anesthetized with an induction dose of 5% isoflurane, followed by a maintenance dose of 2%. Prior to craniotomy surgeries rats received carprofen (5 mg/kg in sterile saline, s.c.) and gentamicin (2.5 mg/kg in sterile saline, i.p.), for analgesic and antibiotic effects, respectively. Guide cannulae (CMA 11 cannulas; CMA, Harvard Apparatus, MA) for microdialysis sampling were implanted at a 10-degree angle in the posterior PVT (pPVT) (-3.0 mm AP; ±1.0 mm ML; -4.5 mm DV, from bregma)^60^. The DV coordinate accounted for the 1-mm length of the microdialysis probe^52^. Cannulae with the wire stylet inserted were secured to the skull with dental cement. One day following surgery rats received an additional dose of carprofen (5 mg/kg s.c.) and were then allowed to recover for at least 5 days before microdialysis experimentation.

### Procedures for habituation prior to in vivo microdialysis sampling (Day 0)

Twenty-four hours before microdialysis collection, rats were temporarily anesthetized with 5% isoflurane using an isoflurane vaporizer, wire stylets were removed, and CMA 11 microdialysis probes (30 kDa cut-off) were inserted unilaterally through the guide cannula to reach the pPVT, secured with a small amount of dental cement, and left in for the duration of the experiment.

Rats were then removed from the vaporizer and allowed to wake up before initiating perfusate into the microdialysis probe. Rats began habituation to the microdialysis sampling chambers between the hours of 8:00 am and 9:00 am. The sampling chambers consisted of a clean cage identical to their homecage (20.32 cm x 21.59 cm x 44.55 cm) and were located inside Med Associates sound-attenuating cabinets (35.56 cm x 56.52 cm x 59.69 cm), which contained a 3-mm circular hole on top for tethered tubing to enter into the chamber. During the habituation session, artificial cerebrospinal fluid (aCSF; 145 mM sodium chloride, 2.68 mM potassium chloride, 1.40 mM calcium chloride, 1.01 mM magnesium sulfate, 1.55 mM dibasic sodium phosphate, 0.45 mM monobasic sodium phosphate, and 0.25 mM ascorbic acid, at a pH=7.4) flowed through the probe for a minimum of 60 min. aCSF mimics the composition of the extracellular fluid, allowing the neurotransmitters present to enter the semipermeable membrane of the probe by passive diffusion. Following the habituation session the probe was subsequently left in the rat brain for all further experimental testing.

### Microdialysis experimental sampling (Day 0 and Day 6)

Experimental dialysate collection occurred on Day 0 prior to the start of PavCA training and then on Day 6 following the completion of PavCA training (Figure 1a). Before dialysate collection on each sampling day, rats were placed into the microdialysis chambers for 120 min, during which probes were continuously perfused with aCSF at a flow rate 2 µL/min for the first hour and 1 µL/min for the second hour. One fraction was collected during each hour, but neither was analyzed. After the 120-min equilibration period, all experimental dialysate samples were collected at 1 µL/min^61^ in 10 min fractions for 90 min, yielding a total of nine 10 µL fractions. These samples were then split into two 5 uL samples: one to be used for neurochemical analysis as described below, and a duplicate for future analysis. Dialysate samples were kept at -80°C until further processing.

### Benzoyl chloride (BzCl) derivatization approach

Prior to high performance liquid chromatography tandem mass spectrometry (HPLC-MS/MS) analysis, neurochemical samples were derivatized. BzCl derivatization is a neurochemical stabilization technique that renders compounds more hydrophobic so they can be separated by reversed phase chromatography. This process increases sensitivity (10-1000-fold) and allows better detection by HPLC-MS/MS^52,62^. BzCl derivatization of dialysates and internal standards were prepared using a modified version of the methods previously published^61^. Briefly, calibration curves were generated using nM standards for all neurochemicals of interest. Samples were derivatized by sequentially adding three reagents to 5μL of dialysate, yielding a sample:buffer:BzCl:internal standard volume ratio of 2:1:1:1, with: 1) 0.1M sodium carbonate buffer (2.5 μL), 2) 2% (v/v) BzCl in acetonitrile (ACN) (2.5µL), and 3) internal standard (489 µL H2O, 489 µL ACN, 10 µL Sulfuric acid and 10 μL Derivatized IS; with concentrations as previously described^61^) (2.5 µL). Following BzCl derivatization, samples were stored at -80°C before HPLC-MS/MS.

### HPLC-MS/MS

Derivatized samples were analyzed using a Phenomenex (Torrance, CA) Kinetex C18 chromatography column (100 × 2.1 mm, 1.7 μm, 100 Å) on a Vanquish Flex ultra high-pressure liquid chromatograph (UHPLC) interfaced to a TSQ Quantum triple quadrupole mass spectrometer (Thermofisher, Waltham, MA). Mobile phase A was 10 mM ammonium formate with 0.1% (v/v) formic acid in water. Mobile phase B was ACN. The following gradient was used: initial, 5% B; 0.05 min, 19% B; 2.00 min, 100% B; 2.25 min, 100% B; 2.30 min, 5% B; 3.2 min, 5% B. The column was held at 30°C in still air mode. Samples were kept at ambient temperature in the autosampler and injected as 5 µL volumes. The spray voltage was set to 3000 V, the vaporizer temperature was set to 300°C, and the auxiliary gas, sheath gas, and ion sweep gas pressures were set at 30, 60, and 0.2 respectively. The capillary temperature was set at 300°C. Positive ion mode was used. Automated peak integration was performed using Thermo X Calibur Quan Browser version 2.1. All peaks were visually inspected to ensure proper integration. Calibration curves were constructed based on peak area ratio (Panalyte/PIS) versus concentrations by linear regression and further analyzed using other metabolomic analysis software (Skyline)^63^. Utilizing a targeted metabolomics approach, a panel of 22 neurobiologically relevant compounds, which include: dopamine (DA), norepinephrine (NE), serotonin (5HT), histamine (Hist), glutamate (Glu), Gamma-aminobutyric acid (GABA), acetylcholine (ACh), choline (Ch), adenosine (Ado),3,4-Dihydroxyphenylacetic acid (DOPAC), 3-Methoxytyramine (3MT), Homovanillic acid (HVA), normetanephrine (NM), taurine (Tau), serine (Ser), aspartate (Asp), glycine (Gly), glutamine (Gln), vanillylmandelic acid (VMA), 3,4-dihydroxyphenylethylene glycol (DOPEG), 3-methoxy-4-hydroxyphenylethylene glycol (MOPEG), and glucose (Glc) were measured via HPLC/MSMS combined with BzCl derivatization methods. The neurochemicals analyzed were further categorized as: Excitatory Neurotransmitters (Glu, Asp); Inhibitory Neurotransmitters and Neuromodulators (GABA, Gly, Ser, Ado, Tau); Cholinergic Compounds (Ch, ACh); Catecholamines and Metabolites (DA, HVA, 3MT, DOPAC, NE, NM, MOPEG, DOPEG, VMA); Biogenic Amine Serotonin (5HT) or Biogenic Amine Histamine (Hist), and Energy Metabolism-Related Compounds (Glc, Gln)^62^.

### Pavlovian conditioned approach (PavCA)

Prior to behavioral testing, rats were given 45-mg banana-flavored pellets (Bioserv; Product# F0059; Flemington, NJ) in a ceramic crock (approximately 30 pellets) in their homecage for 2 days, to familiarize them with the food reward. Pavlovian conditioned approach (PavCA) was conducted as previously described^3^ and occurred in standard behavioral chambers from Med Associates (24.13 cm x 29.21 cm x 30.48 cm) that were encased in sound-attenuating cabinets (35.56 cm x 56.52 cm x 59.69 cm) with ventilation fans that provided air circulation and background noise. A recessed food cup was located in the center of one wall 6 cm above the grid floor. A retractable lever-cue was located to the left or right of the food cup (counterbalanced to control for side bias; 6 cm above the grid floor). The lever-cue was illuminated with an LED backlight upon presentation. A house light was located on the opposite wall 1 cm from the top of the chamber and was illuminated throughout each session.

Behavioral testing for all rats began around 10:30 am each day. Rats were first exposed to one pre-training session consisting of random delivery of 45-mg banana-flavored reward pellets (25 pellets total per session; BioServ; Product#F0059, Flemington, NJ) into the food magazine on a variable interval 30 s schedule (range 0-60 s). The pre-training session lasted approximately 12.5 minutes. Following pre-training, rats underwent 5 sessions of PavCA training. Each session consisted of 25 trials in which the illuminated lever-cue (conditioned stimulus, CS) was presented for 8 s and, upon retraction, a reward pellet (unconditioned stimulus, US) was immediately delivered. Trials were presented on a variable interval 90-s schedule (range 30-150 s). Each PavCA session lasted approximately 40-45 minutes. Similarly, for control animals, all rats received one day of pre-training, followed by 5 sessions of randomized unpaired cue and reward presentations with 25 presentations of each, lasting approximately 40-45 minutes, as previously described^30,31^.

The following measures were collected via Med Associates software for each PavCA session: total number of lever-cue contacts, total number of food cup entries during lever-cue presentation, latency to the first lever-cue contact, and latency to the first food cup entry during lever-cue presentation. These measures allowed us to calculate the PavCA score^41^ for each rat based on the: Response Bias [(lever contacts – food cup entries)/(lever contacts + food cup entries)], Probability Difference [p(lever contact) – p(food cup entry)], and Latency Score [(average response time to enter the food cup – average response time to contact the lever)/duration of the CS presentation]. The average of these three measures was calculated to determine the PavCA score for each of the 5 sessions, which ranged from -1.0 to 1.0. The PavCA index was then generated by averaging the PavCA score from sessions 4 and 5. For this study, a PavCA index ≥0.3 indicates that a rat is a sign-tracker (ST) and interacting more with the lever-cue than food cup upon lever-cue presentation, and a score of <0.3 indicates that a rat is a non-sign-tracker (NST) and either has no preference for the lever-cue versus food cup, or is interacting more with the food cup than the lever-cue upon its presentation.

### Histological assessment of probe placement

Within 24 hours of the final microdialysis collection day, rats were anesthetized with a cocktail of ketamine (90 mg/kg, i.p.) and xylazine (10mg/kg, i.p.). Following injection rats were transcardially perfused with 4% paraformaldehyde. Brains were stored in 4% paraformaldehyde for 24 hours, then moved into 30% sucrose for 3 more days before being cryostat-sectioned into 40 µm coronal cuts. Sections were obtained in 12-well plates, which were then stored in cryoprotectant solution until mounting onto slides. After mounted tissue was completely dry, cresyl-violet staining procedures were used to assess microdialysis probe placement. Criteria for successful placement were defined as visible probe damage below the third ventricle but not surpassing the intermediodorsal nucleus (IMD) (Supporting Figure 1). Microscopy was completed using a brightfield microscope (LEICA DM 1000 LED).

### Software details and statistical approach for data analysis

Statistical analyses were performed using GraphPad Prism version 11.0.0 for Windows (GraphPad Software, Boston, MA, USA). Data are presented as mean ± SEM. Mixed effects models included rat as a random effect to account for repeated measurements within subjects (across training sessions, sampling fractions, and Pre- to Post-time points). Behavioral measures were analyzed across training sessions using mixed effects models, with Session, Phenotype (ST versus NST) and Session x Phenotype interaction as fixed effects (Figure 2). The PavCA index was compared between STs and NSTs using an unpaired t-test (Figure 2). Neurochemical concentrations across a 90-minute experimental sampling period were analyzed within each group (ST, NST, Ctrl) using mixed effects models with Sampling Period, Time Point (Pre-versus Post-PavCA), and their interactions (Figure 3). Neurochemical concentrations averaged across a 90-minute sampling period were analyzed using mixed effects models with Time Point, Group (ST, NST, Ctrl) and Time Point x Group interaction as fixed effects, followed by uncorrected Fisher’s least significant difference (LSD) multiple comparisons where indicated (Figures 3). Data shown in Figure 4 were similarly analyzed, but with Phenotype (ST versus NST) replacing Group, yielding fixed effects of Time Point, Phenotype, and their interaction.

Data shown in Supporting Figure 2 were analyzed using a paired t-test to assess changes in neurochemical concentrations before and after unpaired cue-reward sessions in the control group. Missing values or values below the detection threshold were excluded for the corresponding animal and timepoint. When the estimated between-subject variance was zero, GraphPad Prism excluded the subject random effect from the model. In these cases the denominator degrees of freedom were based on the observation-level residual degrees of freedom rather than the number of subjects. Relationships between Pre-PavCA neurochemical concentrations and PavCA index were assessed using simple linear regression (Figure 3). P-values were adjusted across analytes using the Benjamini–Hochberg procedure (q-value; q<0.05) to account for multiple testing across 22 analytes. Adjusted values are reported in the tables, but interpretation of these exploratory analyses was based largely on unadjusted p-values. Statistical significance was set as p<0.05.

### Software details and statistical approach for multiple linear regression predictive modeling of the PavCA index

All predictive modeling analyses were done using R version 4.5.1. The R package ‘tidyverse’^64^ was used for data processing. Multiple linear regression was fitted using R base command ‘lm’. Stepwise selection using Bayesian information criterion was performed using the ‘step’ command from the ‘stats’ package. The explained sums of squares ratios were computed using ANOVA type II of the ‘car’^65^ package. The model diagnostics were obtained using the ‘performance’^66^ package. All the remaining codes were written using base R functions and commands.

### Data description

Data analysis consisted of n=29 rats (n=14 males, n=15 females) measured at two time points: Pre-PavCA (day 0) and Post-PavCA (day 6). Two phenotypes were derived from the PavCA index as described above (PavCA index <0.3 as NSTs (n=17) and PavCA index ≥ 0.3 as STs (n=12)). Pre-PavCA and Post-PavCA concentrations of 22 analytes were available for each subject. Missing or non-positive values were replaced by the minimum of the available values for the analyte at that time point divided by 5, as imputing values is standard practice in metabolomics data^67,68^. For certain analytes, including ACh, Asp, Ch, DA, NE, NM, Ser, VMA and 3MT some values were missing or labeled as non-positive due to weak analyte signal or concentration falling below the detection threshold, similar to other omics datasets^69-71^. The analytes were grouped into the following 7 categories to aid in interpretation of the results: Excitatory Neurotransmitters (EN): Glu, Asp; Inhibitory Neurotransmitters and Neuromodulators (IN): GABA, Gly, Ser, Ado, Tau; Cholinergic Compounds (CC): Ch, ACh; Catecholamines and Metabolites (CAM): DA, HVA, 3MT, DOPAC, NE, NM, MOPEG, DOPEG, VMA; Biogenic Amine-5HT (BA5): 5HT; Biogenic Amine-Hist (BAH): Hist; Energy Metabolism-Related Compounds (EMRC): Glc, Gln.

### Multiple linear regression modeling

Multiple linear regression (MLR) models were fit to determine whether multivariate PVT neurochemical profiles were associated with the PavCA index. Separate models used either Pre-PavCA or Post-PavCA neurochemical concentrations as candidate predictors, with PavCA index treated as a continuous response variable. Reduced models were selected using stepwise selection based on the Bayesian information criterion (BIC)^72^.

### Model performance measures

Model performance was evaluated using three measures to assess model fit: 1) the coefficient of determination (R^2^), 2) root mean squared error (RMSE), and 3) normalized root mean squared error (NRMSE). Additionally, phenotype-classification accuracy was used to assess the predictive performance of the models. Model-estimated PavCA indices ≥0.3 were classified as STs, whereas values <0.3 were classified as NSTs. Phenotype-classification accuracy was calculated as the proportion of agreement between model-estimated and observed phenotypes. Scatterplots of model-estimated versus observed PavCA indices, color coded by agreement between the model-estimated and observed phenotypes, were generated for each model (Figure 5a, 5d).

### Repeated train-test evaluation

The number of observations for the data set was small (n=29) relative to the number of candidate predictors, increasing the risk of overfitting. Model performance was therefore evaluated using 1000 repeated, phenotype-stratified train-test splits. For each repetition, approximately 20% of STs and 20% of NSTs were randomly assigned to the test set, with the remaining rats assigned to the training set. Stratification ensured that both phenotypes were represented in each test set.

Full models and the BIC-based models were fitted using each training set. The resulting model was then applied to the held-out test set. Test set R^2^, RMSE, NRMSE and phenotype-classification accuracy were calculated for each repetition and summarized across the 1000 repetitions as cross-validated R^2^ (CV-R^2^), CV-RMSE, CV-NRMSE and classification accuracy (CV-accuracy). For CV-R^2^, we considered a value ≥30% to be a moderate fit and ≥50% a good _fit_^73,74^.

### Predictor contribution

The relative contribution of each retained analyte was assessed using a sum of squares ratio (SSRatio). Absolute contribution of each retained analyte was calculated using sum of squares explained by the analyte, defined as the increase in the sum of squared errors (SSE) by excluding the analyte from the model. Then, the SSRatio of the analyte of interest was computed as the percentage of the sum of squares explained by the analyte out of the total of the sum of squares explained by all analytes in the model.

Let *SSE* be the sum of squared errors (SSE) of a model with *p* retained analytes (*p* = 12 for the BIC-based Pre-PavCA model and 14 for the BIC-based post-PavCA model). For a retained analyte *j* (= 1, …, *p*), a reduced model was fit excluding the analyte *j*. Let *SSE*_−j_ be the sum of squared errors of the reduced model. Then, the sum of squares explained by the predictor *j* was computed as the increase in SSE due to excluding analyte *j*, called *Expl*_j_ = *SSE*_−j_ − *SSE*.

### SSRatio of analyte *j* in the model was calculated as

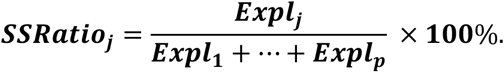

Larger SSRatio values indicate that removal of a predictor produced a greater reduction in model fit. In a model with *p* retained analytes (*p* = 12 for the BIC-based Pre-PavCA model and 14 for the BIC-based post-PavCA model), if all predictors had equal contribution to the model, SSRatio of each predictor would be 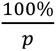. Hence, for a model with *p* retained analytes, 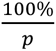 was used as the reference line in the SSRatio plots (Figures 5b and 5e). The bars show how strongly the predictors contributed to the model, compared to this hypothetical equal contribution scenario.

For descriptive purposes, t statistics and corresponding p-values were also obtained from the selected models. P-values were adjusted using the Benjamini-Hochberg procedure and displayed as -log10(adjusted p-value), with a reference line at -log10(0.05) (Figures 5c and 5f). Because the predictors were selected using the same dataset, these post-selection p-values are considered descriptive and not interpreted as confirmatory tests of individual analyte significance.

### Model diagnostics

MLR model assumptions were investigated using standard diagnostic plots and statistics^75,76^. Diagnostics included assessment of 1) linearity, assessed by examining residual plots for systematic nonlinear patterns, 2) homogeneity of variance, assessed by examining residual spread across fitted values, 3) influential observations using Cook’s distance, 4) multicollinearity^77^ using variance inflation factors, and 5) residual normality using quantile-quantile plots.

The model diagnostics for the BIC-based Pre-PavCA and Post-PavCA models did not present any strong evidence of deviations from the model assumptions.

## Acknowledgements

Funding

This research was supported by funding from NIH National Institute on Drug Abuse R01 DA054094 (SBF), R01 DA054094-S1 (SBF, BR), T32 DA 007281 (BR) and F32DA061554 (CNC).

## Abbreviation List

3MT: 3-methoxytyramine
5HT: serotonin
ACh: acetylcholine
aCSF: artificial cerebrospinal fluid
Ado: adenosine
AP: anteroposterior
Asp: aspartate
BA5: biogenic amine (5HT) category
BAH: biogenic amine (histamine) category
BIC: Bayesian information criterion
bHR: bred high-responder
bLR: bred low-responder
BzCl: benzoyl chloride
CAM: catecholamines and metabolites category
CC: cholinergic compounds category
Ch: choline
CS: conditioned stimulus
Ctrl(s): control(s)
CV: cross-validated
DA: dopamine
DOPAC: 3,4-dihydroxyphenylacetic acid
DOPEG: 3,4-dihydroxyphenylethylene glycol
DV: dorsoventral
E: epinephrine
EMRC: energy metabolism-related compounds category
EN: excitatory neurotransmitters category
GABA: γ-aminobutyric acid
Glc: glucose
Gln: glutamine
Glu: glutamate
Gly: glycine
GT(s): goal-tracker(s)
Hist: histamine
HPLC-MS/MS: high-performance liquid chromatography–tandem mass spectrometry
HVA: homovanillic acid
i.p.: intraperitoneal
IMD: intermediodorsal nucleus (of the thalamus)
IN: inhibitory neurotransmitters and neuromodulators category
LC: locus coeruleus
LC-MS/MS: liquid chromatography–tandem mass spectrometry
LH: lateral hypothalamus
ML: mediolateral
MLR: multiple linear regression
MOPEG: 3-methoxy-4-hydroxyphenylethylene glycol
NAc: nucleus accumbens
NE: norepinephrine
NM: normetanephrine
NRMSE: normalized root mean squared error
NST(s): non-sign-tracker(s)
PavCA: Pavlovian conditioned approach
pPVT: posterior paraventricular nucleus of the thalamus
PrL: prelimbic cortex
PVT: paraventricular nucleus of the thalamus
R²: coefficient of determination
RMSE: root mean squared error
s.c.: subcutaneous
Ser: serine
SSE: sum of squared errors
SSRatio: sum of squares ratio
ST(s): sign-tracker(s)
Tau: taurine
TSS: total sum of squares
US: unconditioned stimulus
VIF: variance inflation factor
VMA: vanillylmandelic acid
VTA: ventral tegmental area

## Supporting Information

**Supporting Figure 1.**
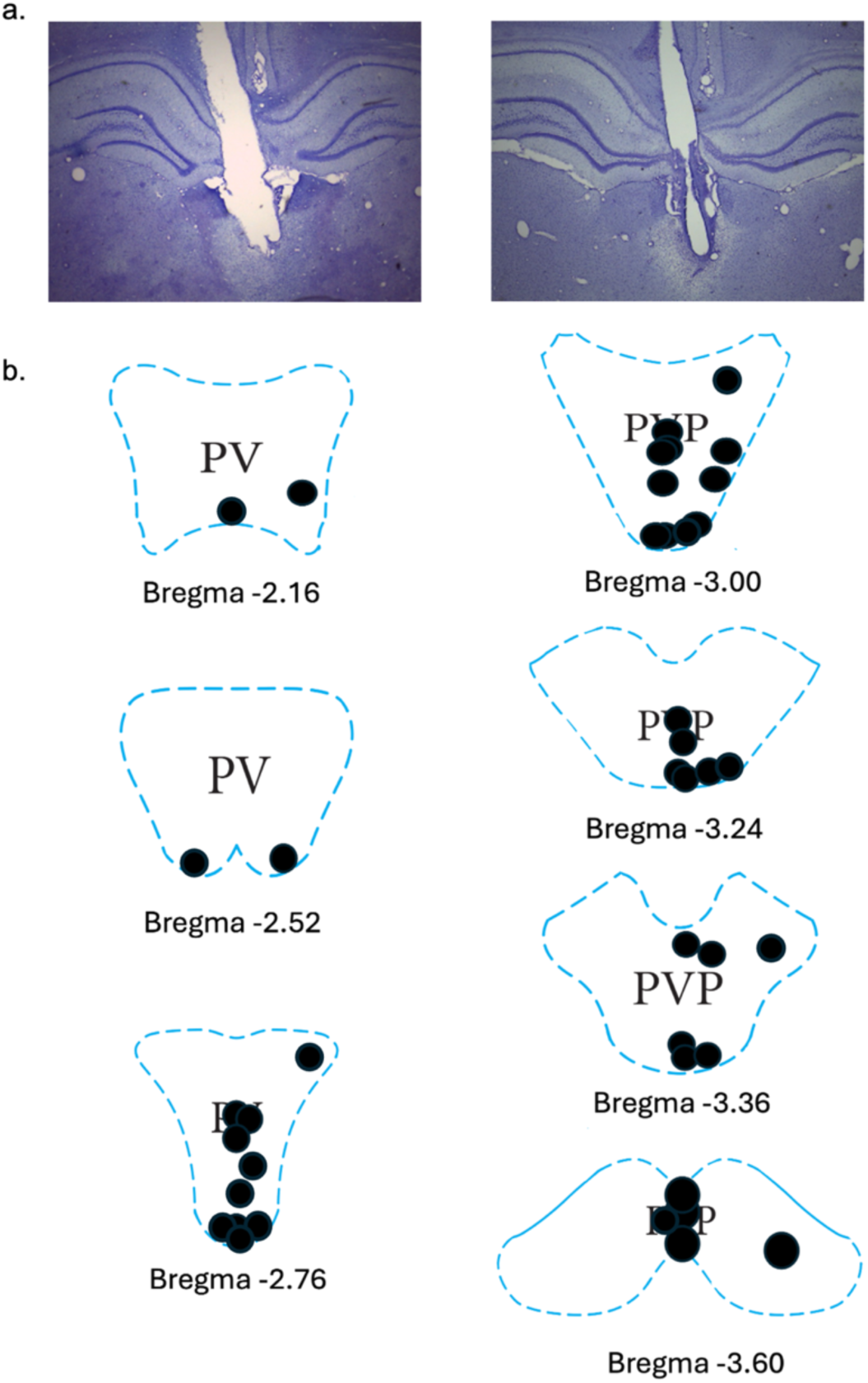
Histology: **(a)** Representative Cresyl-violet stained tissue (40 µm) illustrating microdialysis probe placement in the paraventricular nucleus of the thalamus (PVT) (left image; bregma AP:-2.76 mm) and posterior PVT (right image; bregma AP:-3.24 mm). **(b)** Representative rat brain atlas images of the medial PVT (referred to on atlas as PV) and posterior PVT (referred to on atlas as PVP) indicating stereotaxic coordinates relative to bregma for n=42 subjects. Images are from the George Paxinos and Charles Watson 6th edition with ranges from AP:-2.16 mm to AP:-3.60 mm^60^.

**Supporting Table 1. Analyte Analyses Across 90-min Sampling Period for each Group (ST, NST, Ctrl).** Neurochemical concentrations were analyzed separately within each group using mixed-effects models with Sampling Period (nine 10-min fractions over 90 min), Time Point (Pre versus Post), and their interaction as fixed effects: **(a)** sign-trackers (STs, n=12), **(b)** non-sign-trackers (NSTs, n=17), and **(c)** Controls (Ctrls, n=13), corresponding to the analyses shown in Figure 3a-c. Pre and Post refer to measurements before and after PavCA training for STs and NSTs, or unpaired cue-reward sessions for Ctrls. Reported statistics include the degrees of freedom (DFn and DFd), F-values, p-values, and Benjamini-Hochberg adjusted q-values controlled across the 22 analytes. Unadjusted p-values <0.05 and q-values <0.05 are bolded. For certain analytes, including Asp, Ser, NE, NM, DA and 3MT, some values were missing due to weak signal or concentration falling below the detection threshold.

**Supporting Table 1a:** ST Pre- vs ST Post-PavCA across 90-min sampling period

| Analyte | Fixed effects (type III) | DFn, DFd | F value | p value | q value |
|---|---|---|---|---|---|
| ACh | Sampling Period | 8, 158 | 1.830 | 0.075 | 0.349 |
|  | Time Point | 1, 20 | 2.606 | 0.122 | 0.411 |
|  | Sampling Period x Time Point | 8, 158 | 1.372 | 0.213 | 0.539 |
| Ado | Sampling Period | 8, 173 | 0.878 | 0.537 | 0.847 |
|  | Time Point | 1, 22 | 0.013 | 0.909 | 0.971 |
|  | Sampling Period x Time Point | 8, 173 | 1.016 | 0.426 | 0.847 |
| Asp | Sampling Period | 8, 139 | 0.740 | 0.656 | 0.883 |
|  | Time Point | 1, 18 | 0.015 | 0.903 | 0.971 |
|  | Sampling Period x Time Point | 8, 139 | 0.933 | 0.492 | 0.847 |
| Ch | Sampling Period | 8, 157 | 3.644 | **0.001** | 0.020 |
|  | Time Point | 1, 20 | 0.245 | 0.626 | 0.879 |
|  | Sampling Period x Time Point | 8, 157 | 0.999 | 0.439 | 0.847 |
| DA | Sampling Period | 8, 156 | 1.462 | 0.176 | 0.483 |
|  | Time Point | 1, 20 | 0.047 | 0.831 | 0.941 |
|  | Sampling Period x Time Point | 8, 156 | 2.323 | **0.022** | 0.208 |
| DOPAC | Sampling Period | 8, 173 | 1.873 | 0.067 | 0.349 |
|  | Time Point | 1, 22 | 0.150 | 0.702 | 0.898 |
|  | Sampling Period x Time Point | 8, 173 | 1.611 | 0.125 | 0.411 |
| DOPEG | Sampling Period | 8, 173 | 1.269 | 0.262 | 0.597 |
|  | Time Point | 1, 22 | 9.883 | **0.005** | 0.078 |
|  | Sampling Period x Time Point | 8, 173 | 1.560 | 0.140 | 0.420 |
| GABA | Sampling Period | 8, 173 | 1.855 | 0.070 | 0.349 |
|  | Time Point | 1, 22 | 1.475 | 0.238 | 0.560 |
|  | Sampling Period x Time Point | 8, 173 | 1.945 | 0.056 | 0.349 |
| Gln | Sampling Period | 8, 173 | 2.112 | **0.037** | 0.271 |
|  | Time Point | 1, 22 | 2.638 | 0.119 | 0.411 |
|  | Sampling Period x Time Point | 8, 173 | 0.834 | 0.574 | 0.861 |
| Glu | Sampling Period | 8, 173 | 1.776 | 0.085 | 0.349 |
|  | Time Point | 1, 22 | 0.002 | 0.963 | 0.971 |
|  | Sampling Period x Time Point | 8, 173 | 3.050 | **0.003** | 0.068 |
| Gly | Sampling Period | 8, 172 | 1.909 | 0.062 | 0.349 |
|  | Time Point | 1, 22 | 0.222 | 0.642 | 0.883 |
|  | Sampling Period x Time Point | 8, 172 | 0.341 | 0.949 | 0.971 |
| Hist | Sampling Period | 8, 173 | 0.985 | 0.449 | 0.847 |
|  | Time Point | 1, 22 | 2.762 | 0.111 | 0.411 |
|  | Sampling Period x Time Point | 8, 173 | 0.519 | 0.841 | 0.941 |
| HVA | Sampling Period | 8, 173 | 1.493 | 0.163 | 0.467 |
|  | Time Point | 1, 22 | 0.120 | 0.733 | 0.898 |
|  | Sampling Period x Time Point | 8, 173 | 1.068 | 0.388 | 0.801 |
| MOPEG | Sampling Period | 8, 171 | 0.577 | 0.796 | 0.941 |
|  | Time Point | 1, 22 | 0.774 | 0.388 | 0.801 |
|  | Sampling Period x Time Point | 8, 171 | 0.905 | 0.514 | 0.847 |
| NE | Sampling Period | 8, 173 | 0.870 | 0.543 | 0.847 |
|  | Time Point | 1, 22 | 0.292 | 0.594 | 0.872 |
|  | Sampling Period x Time Point | 8, 173 | 0.872 | 0.542 | 0.847 |
| NM | Sampling Period | 8, 134 | 0.940 | 0.486 | 0.847 |
|  | Time Point | 1, 17 | 6.745 | **0.019** | 0.207 |
|  | Sampling Period x Time Point | 8, 134 | 2.184 | **0.032** | 0.267 |
| Ser | Sampling Period | 8, 154 | 0.677 | 0.711 | 0.898 |
|  | Time Point | 1, 20 | 0.510 | 0.483 | 0.847 |
|  | Sampling Period x Time Point | 8, 154 | 2.519 | **0.013** | 0.176 |
| Tau | Sampling Period | 8, 173 | 1.341 | 0.226 | 0.553 |
|  | Time Point | 1, 22 | 0.001 | 0.971 | 0.971 |
|  | Sampling Period x Time Point | 8, 173 | 0.566 | 0.805 | 0.941 |
| VMA | Sampling Period | 8, 166 | 1.806 | 0.079 | 0.349 |
|  | Time Point | 1, 21 | 0.118 | 0.735 | 0.898 |
|  | Sampling Period x Time Point | 8, 166 | 0.860 | 0.552 | 0.847 |
| 3MT | Sampling Period | 8, 71 | 0.781 | 0.621 | 0.879 |
|  | Time Point | 1, 9 | 0.047 | 0.834 | 0.941 |
|  | Sampling Period x Time Point | 8, 71 | 1.409 | 0.208 | 0.539 |
| 5HT | Sampling Period | 8, 172 | 0.318 | 0.958 | 0.971 |
|  | Time Point | 1, 22 | 0.011 | 0.917 | 0.971 |
|  | Sampling Period x Time Point | 8, 172 | 1.105 | 0.362 | 0.796 |

**Supporting Table 1b:**
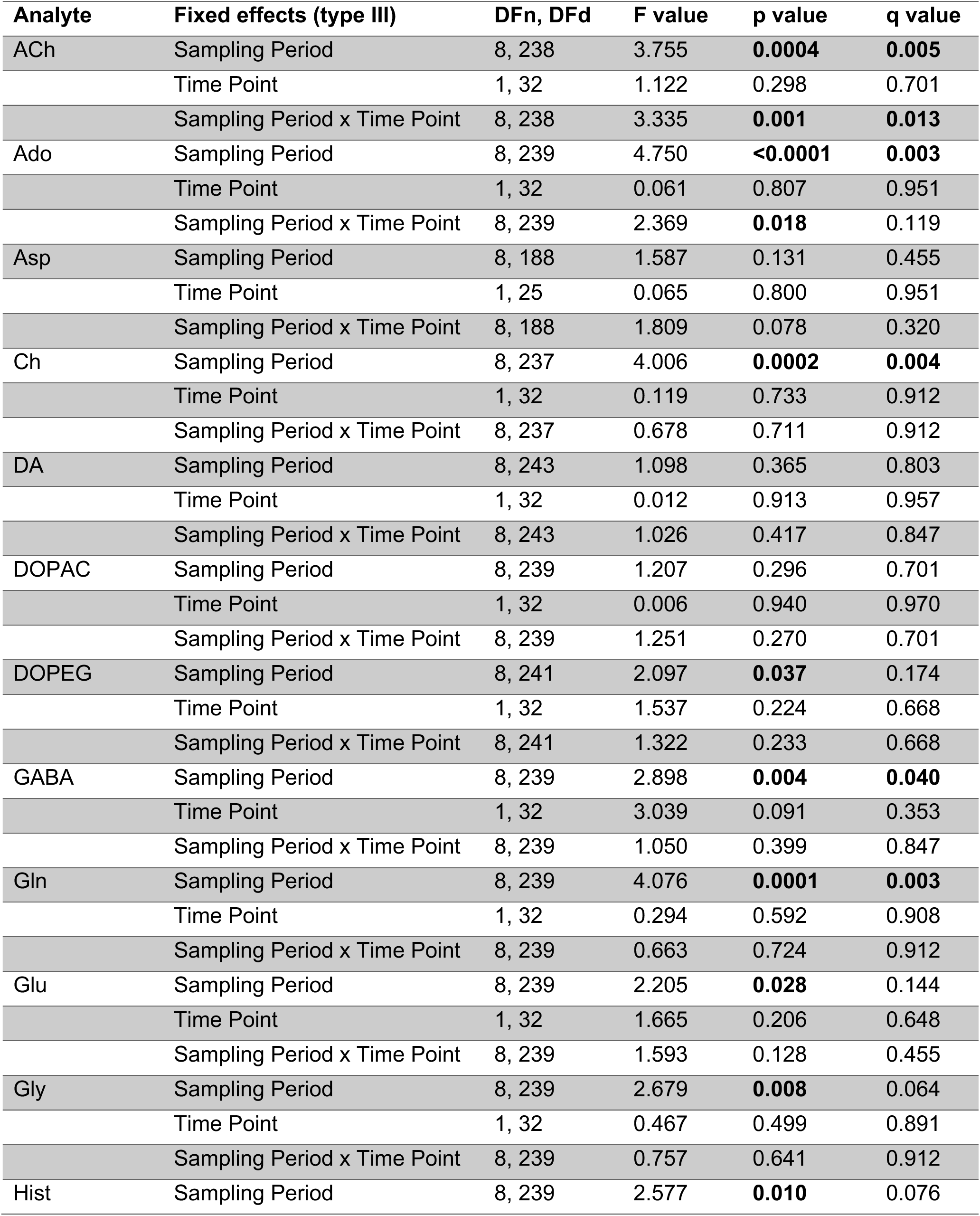
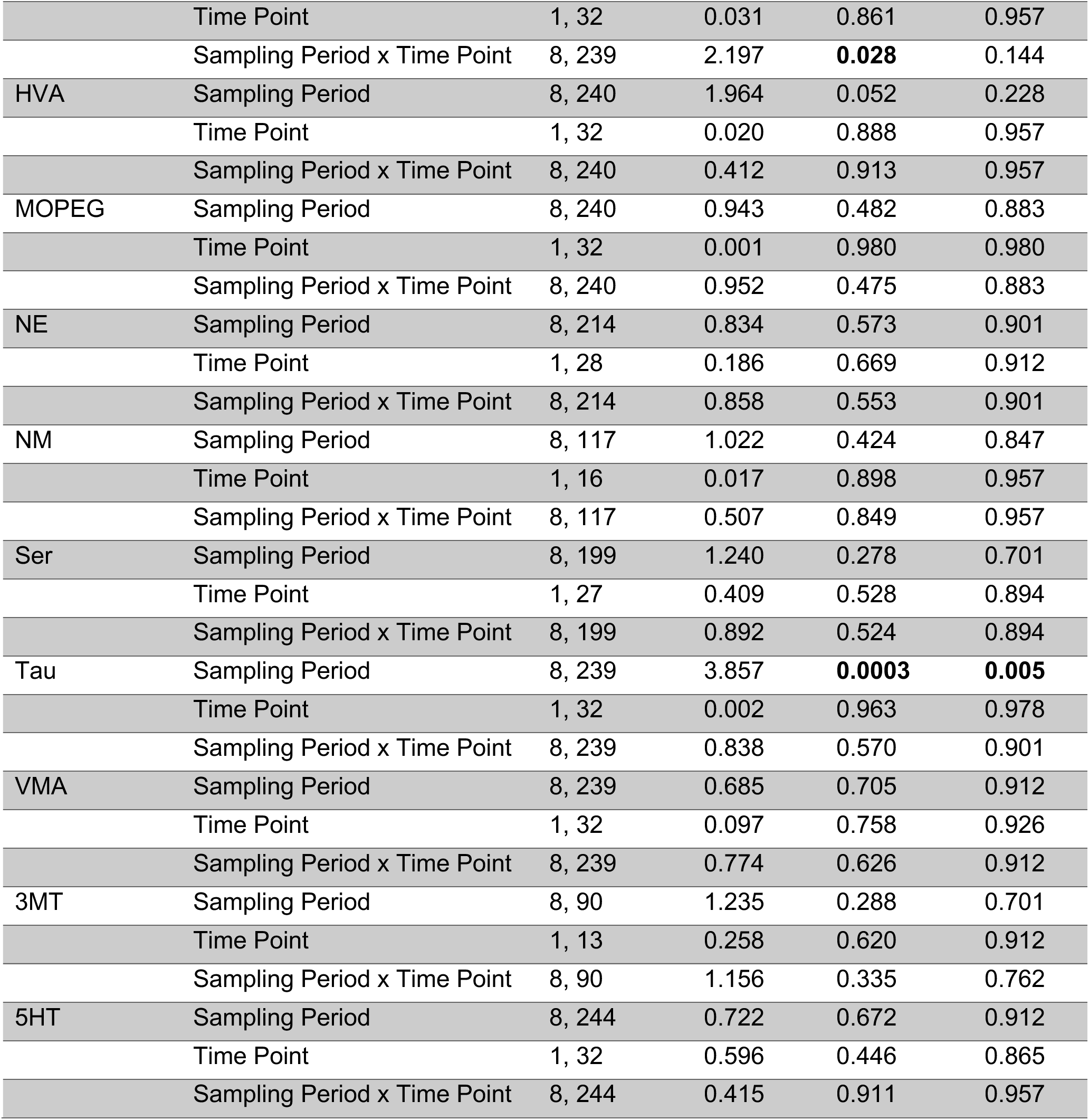
NST Pre- vs NST Post-PavCA across 90-min sampling period

| Analyte | Fixed effects (type III) | DFn, DFd | F value | p value | q value |
|---|---|---|---|---|---|
| ACh | Sampling Period | 8, 238 | 3.755 | **0.0004** | **0.005** |
|  | Time Point | 1, 32 | 1.122 | 0.298 | 0.701 |
|  | Sampling Period x Time Point | 8, 238 | 3.335 | **0.001** | **0.013** |
| Ado | Sampling Period | 8, 239 | 4.750 | **<0.0001** | **0.003** |
|  | Time Point | 1, 32 | 0.061 | 0.807 | 0.951 |
|  | Sampling Period x Time Point | 8, 239 | 2.369 | **0.018** | 0.119 |
| Asp | Sampling Period | 8, 188 | 1.587 | 0.131 | 0.455 |
|  | Time Point | 1, 25 | 0.065 | 0.800 | 0.951 |
|  | Sampling Period x Time Point | 8, 188 | 1.809 | 0.078 | 0.320 |
| Ch | Sampling Period | 8, 237 | 4.006 | **0.0002** | **0.004** |
|  | Time Point | 1, 32 | 0.119 | 0.733 | 0.912 |
|  | Sampling Period x Time Point | 8, 237 | 0.678 | 0.711 | 0.912 |
| DA | Sampling Period | 8, 243 | 1.098 | 0.365 | 0.803 |
|  | Time Point | 1, 32 | 0.012 | 0.913 | 0.957 |
|  | Sampling Period x Time Point | 8, 243 | 1.026 | 0.417 | 0.847 |
| DOPAC | Sampling Period | 8, 239 | 1.207 | 0.296 | 0.701 |
|  | Time Point | 1, 32 | 0.006 | 0.940 | 0.970 |
|  | Sampling Period x Time Point | 8, 239 | 1.251 | 0.270 | 0.701 |
| DOPEG | Sampling Period | 8, 241 | 2.097 | **0.037** | 0.174 |
|  | Time Point | 1, 32 | 1.537 | 0.224 | 0.668 |
|  | Sampling Period x Time Point | 8, 241 | 1.322 | 0.233 | 0.668 |
| GABA | Sampling Period | 8, 239 | 2.898 | **0.004** | **0.040** |
|  | Time Point | 1, 32 | 3.039 | 0.091 | 0.353 |
|  | Sampling Period x Time Point | 8, 239 | 1.050 | 0.399 | 0.847 |
| Gln | Sampling Period | 8, 239 | 4.076 | **0.0001** | **0.003** |
|  | Time Point | 1, 32 | 0.294 | 0.592 | 0.908 |
|  | Sampling Period x Time Point | 8, 239 | 0.663 | 0.724 | 0.912 |
| Glu | Sampling Period | 8, 239 | 2.205 | **0.028** | 0.144 |
|  | Time Point | 1, 32 | 1.665 | 0.206 | 0.648 |
|  | Sampling Period x Time Point | 8, 239 | 1.593 | 0.128 | 0.455 |
| Gly | Sampling Period | 8, 239 | 2.679 | **0.008** | 0.064 |
|  | Time Point | 1, 32 | 0.467 | 0.499 | 0.891 |
|  | Sampling Period x Time Point | 8, 239 | 0.757 | 0.641 | 0.912 |
| Hist | Sampling Period | 8, 239 | 2.577 | **0.010** | 0.076 |
|  | Time Point | 1, 32 | 0.031 | 0.861 | 0.957 |
|  | Sampling Period x Time Point | 8, 239 | 2.197 | **0.028** | 0.144 |
| HVA | Sampling Period | 8, 240 | 1.964 | 0.052 | 0.228 |
|  | Time Point | 1, 32 | 0.020 | 0.888 | 0.957 |
|  | Sampling Period x Time Point | 8, 240 | 0.412 | 0.913 | 0.957 |
| MOPEG | Sampling Period | 8, 240 | 0.943 | 0.482 | 0.883 |
|  | Time Point | 1, 32 | 0.001 | 0.980 | 0.980 |
|  | Sampling Period x Time Point | 8, 240 | 0.952 | 0.475 | 0.883 |
| NE | Sampling Period | 8, 214 | 0.834 | 0.573 | 0.901 |
|  | Time Point | 1, 28 | 0.186 | 0.669 | 0.912 |
|  | Sampling Period x Time Point | 8, 214 | 0.858 | 0.553 | 0.901 |
| NM | Sampling Period | 8, 117 | 1.022 | 0.424 | 0.847 |
|  | Time Point | 1, 16 | 0.017 | 0.898 | 0.957 |
|  | Sampling Period x Time Point | 8, 117 | 0.507 | 0.849 | 0.957 |
| Ser | Sampling Period | 8, 199 | 1.240 | 0.278 | 0.701 |
|  | Time Point | 1, 27 | 0.409 | 0.528 | 0.894 |
|  | Sampling Period x Time Point | 8, 199 | 0.892 | 0.524 | 0.894 |
| Tau | Sampling Period | 8, 239 | 3.857 | **0.0003** | **0.005** |
|  | Time Point | 1, 32 | 0.002 | 0.963 | 0.978 |
|  | Sampling Period x Time Point | 8, 239 | 0.838 | 0.570 | 0.901 |
| VMA | Sampling Period | 8, 239 | 0.685 | 0.705 | 0.912 |
|  | Time Point | 1, 32 | 0.097 | 0.758 | 0.926 |
|  | Sampling Period x Time Point | 8, 239 | 0.774 | 0.626 | 0.912 |
| 3MT | Sampling Period | 8, 90 | 1.235 | 0.288 | 0.701 |
|  | Time Point | 1, 13 | 0.258 | 0.620 | 0.912 |
|  | Sampling Period x Time Point | 8, 90 | 1.156 | 0.335 | 0.762 |
| 5HT | Sampling Period | 8, 244 | 0.722 | 0.672 | 0.912 |
|  | Time Point | 1, 32 | 0.596 | 0.446 | 0.865 |
|  | Sampling Period x Time Point | 8, 244 | 0.415 | 0.911 | 0.957 |

**Supporting Table 1c:**
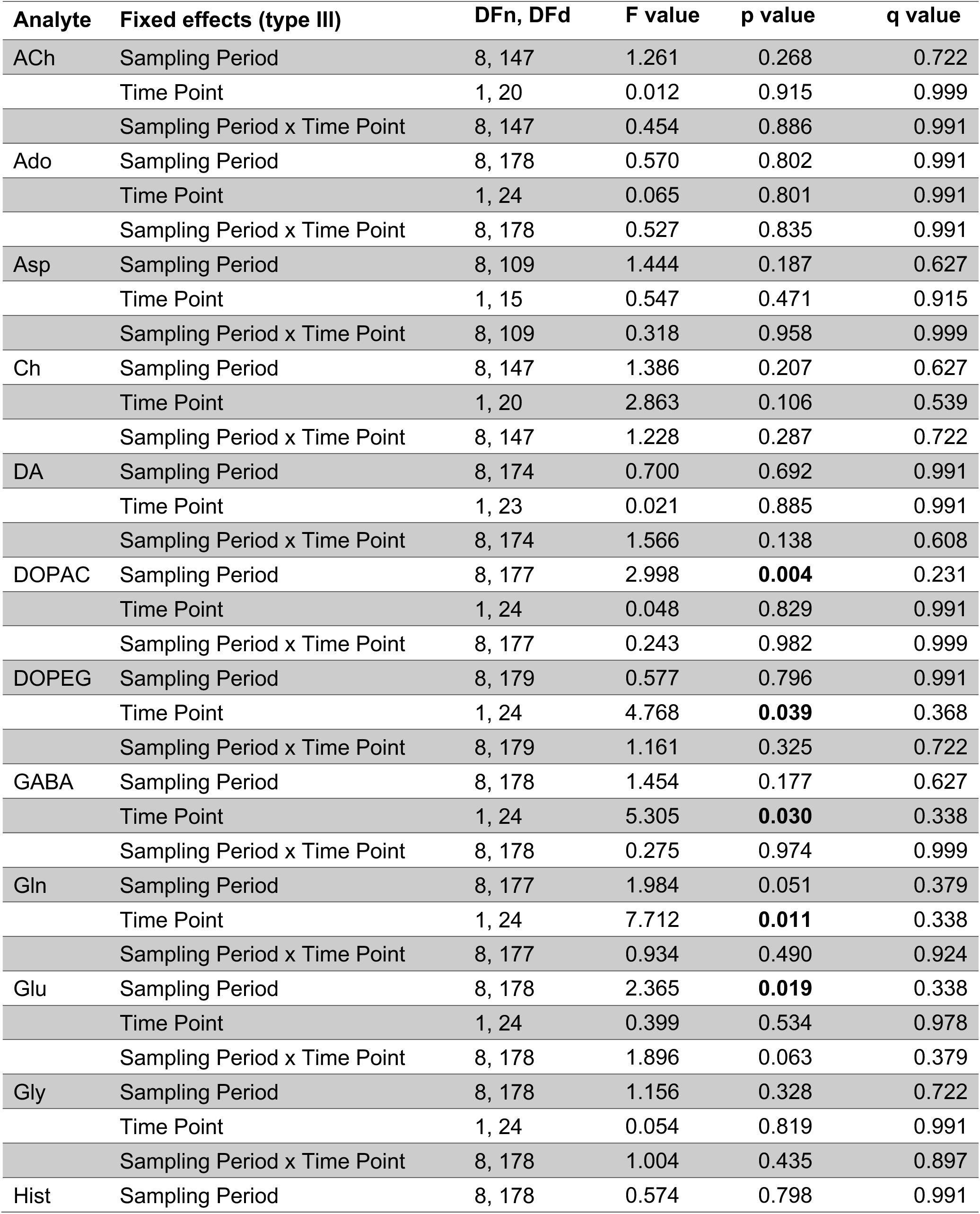
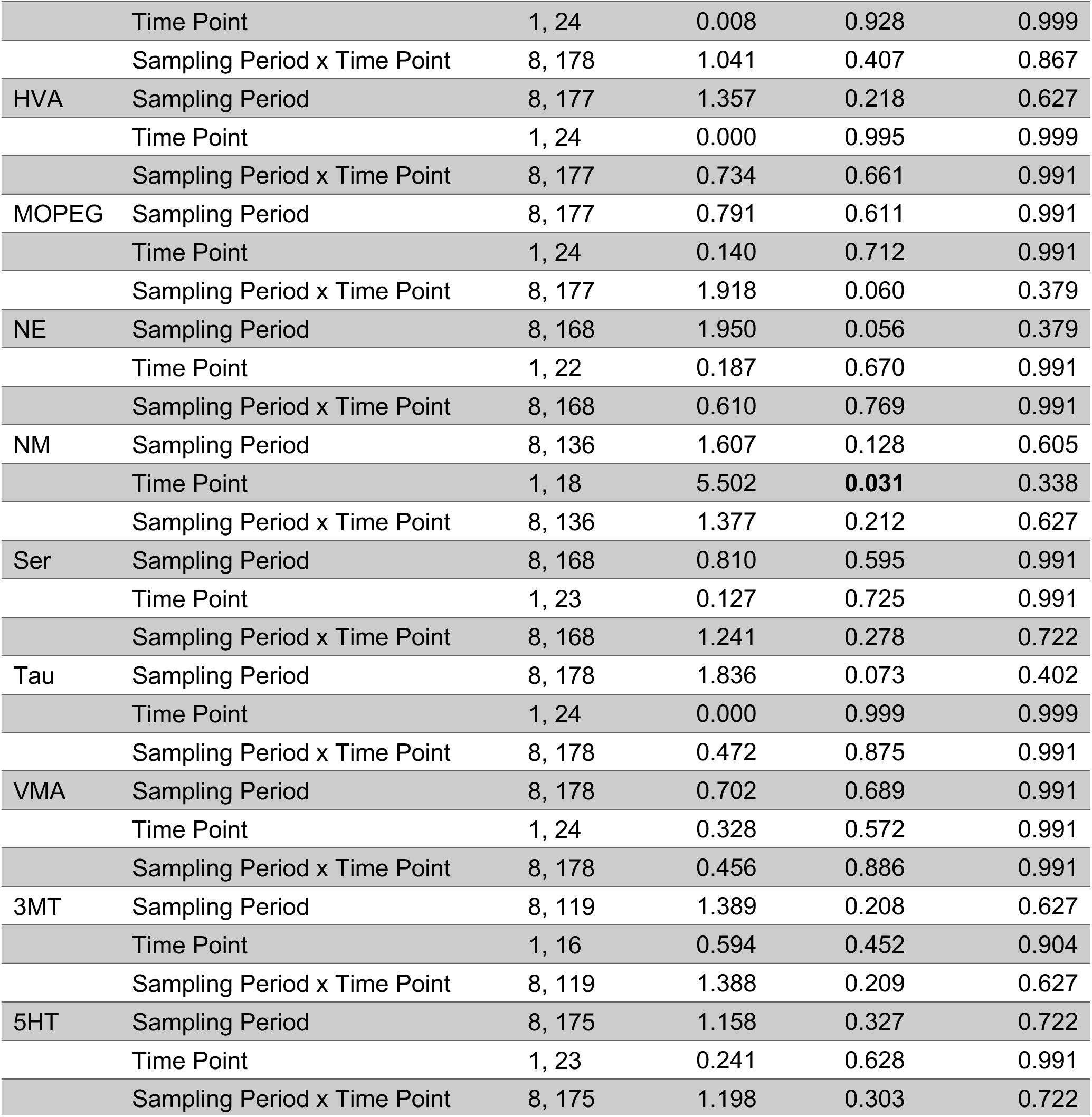
Ctrl Pre- vs Ctrl Post-Unpaired Sessions across 90-min sampling period

| Analyte | Fixed effects (type III) | DFn, DFd | F value | p value | q value |
|---|---|---|---|---|---|
| ACh | Sampling Period | 8, 147 | 1.261 | 0.268 | 0.722 |
|  | Time Point | 1, 20 | 0.012 | 0.915 | 0.999 |
|  | Sampling Period x Time Point | 8, 147 | 0.454 | 0.886 | 0.991 |
| Ado | Sampling Period | 8, 178 | 0.570 | 0.802 | 0.991 |
|  | Time Point | 1, 24 | 0.065 | 0.801 | 0.991 |
|  | Sampling Period x Time Point | 8, 178 | 0.527 | 0.835 | 0.991 |
| Asp | Sampling Period | 8, 109 | 1.444 | 0.187 | 0.627 |
|  | Time Point | 1, 15 | 0.547 | 0.471 | 0.915 |
|  | Sampling Period x Time Point | 8, 109 | 0.318 | 0.958 | 0.999 |
| Ch | Sampling Period | 8, 147 | 1.386 | 0.207 | 0.627 |
|  | Time Point | 1, 20 | 2.863 | 0.106 | 0.539 |
|  | Sampling Period x Time Point | 8, 147 | 1.228 | 0.287 | 0.722 |
| DA | Sampling Period | 8, 174 | 0.700 | 0.692 | 0.991 |
|  | Time Point | 1, 23 | 0.021 | 0.885 | 0.991 |
|  | Sampling Period x Time Point | 8, 174 | 1.566 | 0.138 | 0.608 |
| DOPAC | Sampling Period | 8, 177 | 2.998 | **0.004** | 0.231 |
|  | Time Point | 1, 24 | 0.048 | 0.829 | 0.991 |
|  | Sampling Period x Time Point | 8, 177 | 0.243 | 0.982 | 0.999 |
| DOPEG | Sampling Period | 8, 179 | 0.577 | 0.796 | 0.991 |
|  | Time Point | 1, 24 | 4.768 | **0.039** | 0.368 |
|  | Sampling Period x Time Point | 8, 179 | 1.161 | 0.325 | 0.722 |
| GABA | Sampling Period | 8, 178 | 1.454 | 0.177 | 0.627 |
|  | Time Point | 1, 24 | 5.305 | **0.030** | 0.338 |
|  | Sampling Period x Time Point | 8, 178 | 0.275 | 0.974 | 0.999 |
| Gln | Sampling Period | 8, 177 | 1.984 | 0.051 | 0.379 |
|  | Time Point | 1, 24 | 7.712 | **0.011** | 0.338 |
|  | Sampling Period x Time Point | 8, 177 | 0.934 | 0.490 | 0.924 |
| Glu | Sampling Period | 8, 178 | 2.365 | **0.019** | 0.338 |
|  | Time Point | 1, 24 | 0.399 | 0.534 | 0.978 |
|  | Sampling Period x Time Point | 8, 178 | 1.896 | 0.063 | 0.379 |
| Gly | Sampling Period | 8, 178 | 1.156 | 0.328 | 0.722 |
|  | Time Point | 1, 24 | 0.054 | 0.819 | 0.991 |
|  | Sampling Period x Time Point | 8, 178 | 1.004 | 0.435 | 0.897 |
| Hist | Sampling Period | 8, 178 | 0.574 | 0.798 | 0.991 |
|  | Time Point | 1, 24 | 0.008 | 0.928 | 0.999 |
|  | Sampling Period x Time Point | 8, 178 | 1.041 | 0.407 | 0.867 |
| HVA | Sampling Period | 8, 177 | 1.357 | 0.218 | 0.627 |
|  | Time Point | 1, 24 | 0.000 | 0.995 | 0.999 |
|  | Sampling Period x Time Point | 8, 177 | 0.734 | 0.661 | 0.991 |
| MOPEG | Sampling Period | 8, 177 | 0.791 | 0.611 | 0.991 |
|  | Time Point | 1, 24 | 0.140 | 0.712 | 0.991 |
|  | Sampling Period x Time Point | 8, 177 | 1.918 | 0.060 | 0.379 |
| NE | Sampling Period | 8, 168 | 1.950 | 0.056 | 0.379 |
|  | Time Point | 1, 22 | 0.187 | 0.670 | 0.991 |
|  | Sampling Period x Time Point | 8, 168 | 0.610 | 0.769 | 0.991 |
| NM | Sampling Period | 8, 136 | 1.607 | 0.128 | 0.605 |
|  | Time Point | 1, 18 | 5.502 | **0.031** | 0.338 |
|  | Sampling Period x Time Point | 8, 136 | 1.377 | 0.212 | 0.627 |
| Ser | Sampling Period | 8, 168 | 0.810 | 0.595 | 0.991 |
|  | Time Point | 1, 23 | 0.127 | 0.725 | 0.991 |
|  | Sampling Period x Time Point | 8, 168 | 1.241 | 0.278 | 0.722 |
| Tau | Sampling Period | 8, 178 | 1.836 | 0.073 | 0.402 |
|  | Time Point | 1, 24 | 0.000 | 0.999 | 0.999 |
|  | Sampling Period x Time Point | 8, 178 | 0.472 | 0.875 | 0.991 |
| VMA | Sampling Period | 8, 178 | 0.702 | 0.689 | 0.991 |
|  | Time Point | 1, 24 | 0.328 | 0.572 | 0.991 |
|  | Sampling Period x Time Point | 8, 178 | 0.456 | 0.886 | 0.991 |
| 3MT | Sampling Period | 8, 119 | 1.389 | 0.208 | 0.627 |
|  | Time Point | 1, 16 | 0.594 | 0.452 | 0.904 |
|  | Sampling Period x Time Point | 8, 119 | 1.388 | 0.209 | 0.627 |
| 5HT | Sampling Period | 8, 175 | 1.158 | 0.327 | 0.722 |
|  | Time Point | 1, 23 | 0.241 | 0.628 | 0.991 |
|  | Sampling Period x Time Point | 8, 175 | 1.198 | 0.303 | 0.722 |

**Supporting Table 2.** Analyses of mean neurochemical concentrations by Time Point and Group (ST, NST, Ctrl). Mean neurochemical concentrations averaged across the 90-min sampling period were analyzed using mixed-effects models with Time Point (Pre versus Post), Group (ST (n=12), NST (n=17), Ctrl (n=13)), and their interaction as fixed effects. Pre and Post refer to measurements before and after PavCA training for STs and NSTs, or unpaired cue-reward sessions for Ctrls. These analyses correspond to Figure 3d and include 21 additional analytes. Reported statistics include numerator and denominator degrees of freedom (DFn, DFd), F-values, p-values, and Benjamini–Hochberg-adjusted q-values controlled across the 22 analytes. Unadjusted p-values <0.05 and q-values <0.05 are bolded. For certain analytes, including Asp, Ser, NE, NM, DA and 3MT, some values were missing due to weak signal or concentration falling below the detection threshold.

| Analyte | Fixed effects (type III) | DFn, DFd | F value | p value | q value |
|---|---|---|---|---|---|
| ACh | Time Point | 1, 36 | 1.919 | 0.175 | 0.576 |
|  | Group | 2, 36 | 0.266 | 0.768 | 0.859 |
|  | Time Point x Group | 2, 36 | 0.556 | 0.579 | 0.759 |
| Ado | Time Point | 1, 39 | 0.498 | 0.485 | 0.748 |
|  | Group | 2, 39 | 2.224 | 0.122 | 0.510 |
|  | Time Point x Group | 2, 39 | 0.611 | 0.548 | 0.754 |
| Asp | Time Point | 1, 23 | 0.933 | 0.344 | 0.727 |
|  | Group | 2, 37 | 0.699 | 0.503 | 0.748 |
|  | Time Point x Group | 2, 23 | 0.801 | 0.461 | 0.748 |
| Ch | Time Point | 1, 36 | 3.011 | 0.091 | 0.463 |
|  | Group | 2, 36 | 0.728 | 0.490 | 0.748 |
|  | Time Point x Group | 2, 36 | 0.895 | 0.418 | 0.748 |
| DA | Time Point | 1, 36 | 1.262 | 0.269 | 0.723 |
|  | Group | 2, 39 | 2.980 | 0.063 | 0.375 |
|  | Time Point x Group | 2, 36 | 0.522 | 0.598 | 0.759 |
| DOPAC | Time Point | 1, 38 | 0.997 | 0.324 | 0.727 |
|  | Group | 2, 39 | 0.666 | 0.520 | 0.748 |
|  | Time Point x Group | 2, 38 | 0.468 | 0.630 | 0.770 |
| DOPEG | Time Point | 1, 39 | 40.990 | **<0.0001** | **0.003** |
|  | Group | 2, 39 | 0.285 | 0.754 | 0.859 |
|  | Time Point x Group | 2, 39 | 2.981 | 0.062 | 0.375 |
| GABA | Time Point | 1, 39 | 14.740 | **0.0004** | **0.009** |
|  | Group | 2, 39 | 0.216 | 0.807 | 0.859 |
|  | Time Point x Group | 2, 39 | 0.648 | 0.529 | 0.748 |
| Gln | Time Point | 1, 39 | 13.790 | **0.001** | **0.010** |
|  | Group | 2, 39 | 1.188 | 0.316 | 0.727 |
|  | Time Point x Group | 2, 39 | 2.185 | 0.126 | 0.510 |
| Glu | Time Point | 1, 39 | 0.842 | 0.365 | 0.727 |
|  | Group | 2, 39 | 0.444 | 0.645 | 0.774 |
|  | Time Point x Group | 2, 39 | 0.244 | 0.785 | 0.859 |
| Gly | Time Point | 1, 39 | 0.969 | 0.331 | 0.727 |
|  | Group | 2, 39 | 1.340 | 0.274 | 0.723 |
|  | Time Point x Group | 2, 39 | 0.061 | 0.941 | 0.963 |
| Hist | Time Point | 1, 39 | 4.468 | **0.041** | 0.338 |
|  | Group | 2, 39 | 1.907 | 0.162 | 0.563 |
|  | Time Point x Group | 2, 39 | 3.211 | 0.051 | 0.375 |
| HVA | Time Point | 1, 39 | 0.770 | 0.386 | 0.727 |
|  | Group | 2, 39 | 0.788 | 0.462 | 0.748 |
|  | Time Point x Group | 2, 39 | 0.321 | 0.728 | 0.857 |
| MOPEG | Time Point | 1, 39 | 1.721 | 0.197 | 0.620 |
|  | Group | 2, 39 | 0.641 | 0.532 | 0.748 |
|  | Time Point x Group | 2, 39 | 0.993 | 0.380 | 0.727 |
| NE | Time Point | 1, 72 | 0.905 | 0.345 | 0.727 |
|  | Group | 2, 72 | 1.045 | 0.357 | 0.727 |
|  | Time Point x Group | 2, 72 | 0.688 | 0.506 | 0.748 |
| NM | Time Point | 1, 20 | 13.340 | **0.002** | **0.021** |
|  | Group | 2, 30 | 3.636 | **0.039** | 0.338 |
|  | Time Point x Group | 2, 20 | 2.410 | 0.115 | 0.510 |
| Ser | Time Point | 1, 31 | 0.036 | 0.850 | 0.891 |
|  | Group | 2, 39 | 1.402 | 0.258 | 0.723 |
|  | Time Point x Group | 2, 31 | 0.883 | 0.424 | 0.748 |
| Tau | Time Point | 1, 39 | 0.065 | 0.801 | 0.859 |
|  | Group | 2, 39 | 2.042 | 0.144 | 0.526 |
|  | Time Point x Group | 2, 39 | 0.053 | 0.949 | 0.963 |
| VMA | Time Point | 1, 38 | 0.819 | 0.371 | 0.727 |
|  | Group | 2, 39 | 0.526 | 0.595 | 0.759 |
|  | Time Point x Group | 2, 38 | 2.731 | 0.078 | 0.429 |
| 3MT | Time Point | 1, 12 | 0.064 | 0.804 | 0.859 |
|  | Group | 2, 26 | 2.197 | 0.131 | 0.510 |
|  | Time Point x Group | 2, 12 | 0.849 | 0.452 | 0.748 |
| 5HT | Time Point | 1, 36 | 1.480 | 0.232 | 0.695 |
|  | Group | 2, 38 | 0.493 | 0.614 | 0.765 |
|  | Time Point x Group | 2, 36 | 0.021 | 0.979 | 0.979 |

**Supporting Table 3.** Simple linear regression analyses of associations between Pre-PavCA analyte concentrations and subsequent PavCA index. Simple linear regression was used to examine associations between Pre-PavCA analyte concentrations and subsequent PavCA index, corresponding to the analysis shown in Figure 3e for the additional analytes. Reported statistics include numerator and denominator degrees of freedom (DFn, DFd), F-values, p-values, Benjamini–Hochberg-adjusted q-values, and R² values. Unadjusted p-values <0.05 and q-values <0.05 are bolded. For certain analytes, including ACh, Asp, Ch, DA, NE, NM, Ser, VMA and 3MT some values were missing due to weak signal or concentration falling below the detection threshold.

| **Analyte** | **DFn, DFd** | **F value** | **p value** | **q value** | **R<sup>2</sup> value** |
|---|---|---|---|---|---|
| ACh | 1, 26 | 0.148 | 0.703 | 0.737 | 0.006 |
| Ado | 1, 27 | 3.831 | 0.061 | 0.372 | 0.124 |
| Asp | 1, 23 | 0.330 | 0.572 | 0.737 | 0.014 |
| Ch | 1, 26 | 0.070 | 0.793 | 0.793 | 0.003 |
| DA | 1, 26 | 0.805 | 0.378 | 0.737 | 0.030 |
| DOPAC | 1, 27 | 0.475 | 0.497 | 0.737 | 0.017 |
| DOPEG | 1, 27 | 3.025 | 0.093 | 0.411 | 0.101 |
| GABA | 1, 27 | 0.248 | 0.623 | 0.737 | 0.009 |
| Glc | 1, 27 | 6.091 | 0.020 | 0.372 | 0.184 |
| Gln | 1, 27 | 0.352 | 0.558 | 0.737 | 0.013 |
| Glu | 1, 27 | 0.159 | 0.693 | 0.737 | 0.006 |
| Gly | 1, 27 | 0.484 | 0.493 | 0.737 | 0.018 |
| Hist | 1, 27 | 3.88 | 0.059 | 0.372 | 0.126 |
| HVA | 1, 27 | 0.247 | 0.624 | 0.737 | 0.009 |
| MOPEG | 1, 27 | 0.822 | 0.373 | 0.737 | 0.030 |
| NE | 1, 26 | 0.256 | 0.617 | 0.737 | 0.010 |
| NM | 1, 19 | 2.415 | 0.137 | 0.430 | 0.113 |
| Ser | 1, 26 | 1.916 | 0.178 | 0.490 | 0.069 |
| Tau | 1, 27 | 0.770 | 0.388 | 0.737 | 0.028 |
| VMA | 1, 27 | 0.148 | 0.704 | 0.737 | 0.005 |
| 3MT | 1, 13 | 3.973 | 0.068 | 0.372 | 0.234 |
| 5HT | 1, 27 | 2.679 | 0.113 | 0.415 | 0.090 |

**Supporting Table 4.**
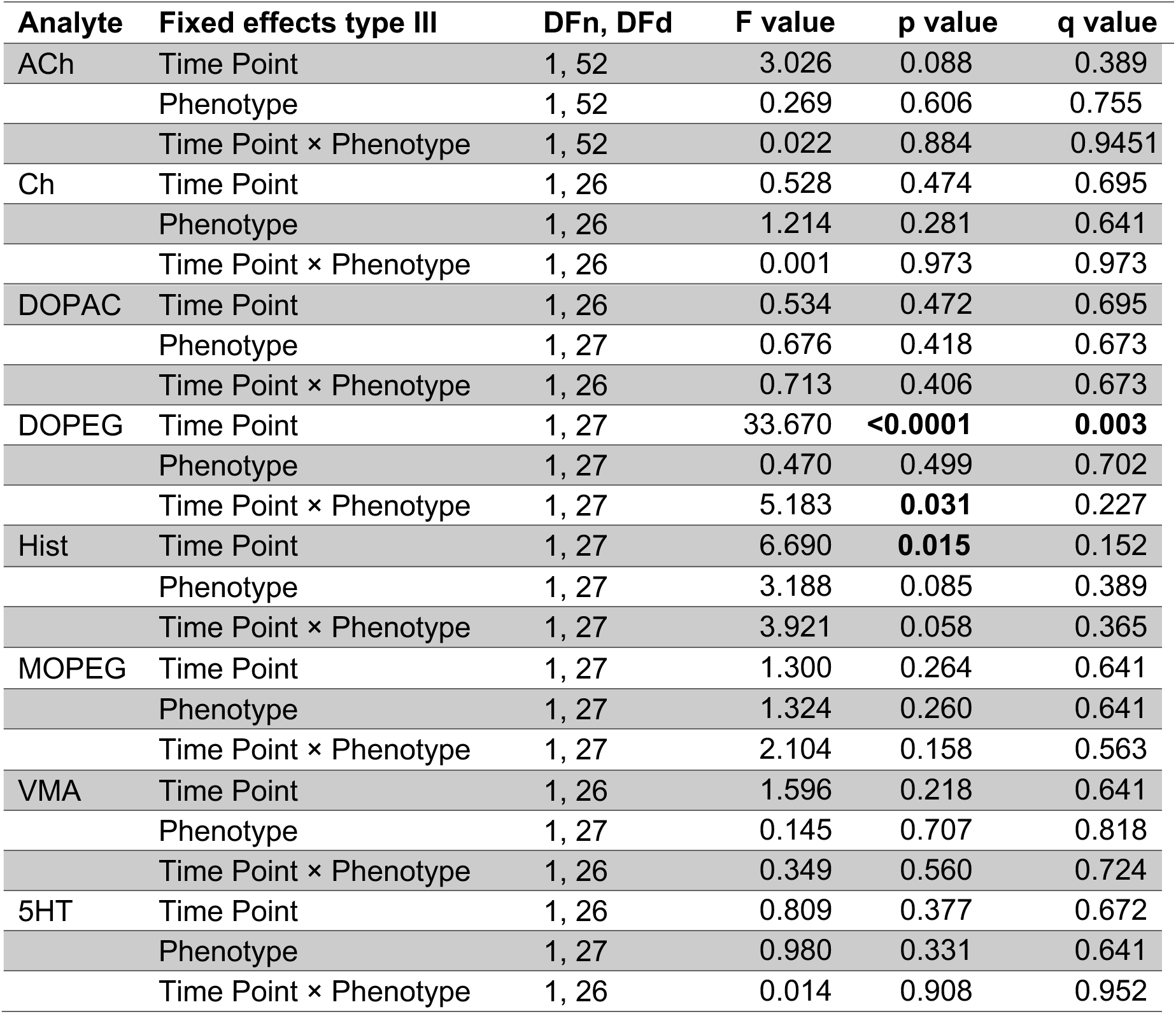
Analyses of mean neurochemical concentrations by Time Point and Phenotype (ST versus NST). Mean neurochemical concentrations averaged across the 90-min sampling period were analyzed using mixed-effects models with Time Point (Pre-versus Post-PavCA), Phenotype (ST versus NST), and their interaction as fixed effects. These analyses correspond to those shown in Figure 4a–n and include additional analytes. Reported statistics include numerator and denominator degrees of freedom (DFn, DFd), F-values, p-values, and Benjamini–Hochberg-adjusted q-values (Table 3). Unadjusted p-values <0.05 and q-values <0.05 are bolded. For certain analytes, including ACh, Ch, and VMA, some values were missing due to weak signal or concentration falling below the detection threshold.

**Supporting Figure 2.**
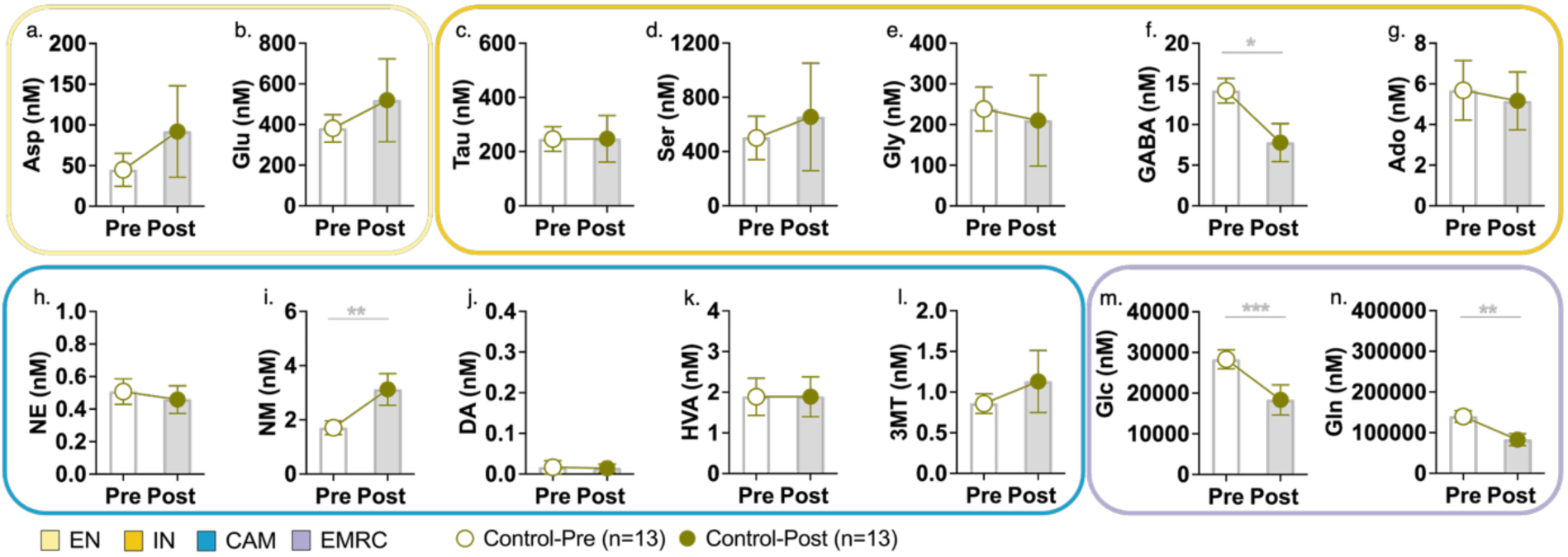
PVT neurochemical concentrations before and after unpaired cue-reward sessions in control (Ctrl) rats. **(a–n)** Mean ± SEM concentrations (nM) averaged across the 90-min sampling period are shown before (Pre, white bars) and after (Post, gray bars) unpaired sessions (n=13; analyte-specific sample sizes vary due to missing values). Paired t-tests identified significant Pre–Post differences for **(f)** γ-aminobutyric acid (GABA; *p=0.027), **(i)** normetanephrine (NM; **p=0.005), **(m)** glucose (Glc; ***p=0.0006), and **(n)** glutamine (Gln; **p=0.002). No significant differences were detected for **(a)** aspartate (Asp), **(b)** glutamate (Glu), **(c)** taurine (Tau), **(d)** serine (Ser), **(e)** glycine (Gly), **(g)** adenosine (Ado), **(h)** norepinephrine (NE), **(j)** dopamine (DA), **(k)** 3-methoxytyramine (3MT), or **(l)** homovanillic acid (HVA). Significance annotations reflect unadjusted p-values. Neurochemical categories are excitatory neurotransmitters (EN, light yellow), inhibitory neurotransmitters and neuromodulators (IN, dark yellow), catecholamines and metabolites (CAM, cerulean blue), and energy metabolism-related compounds (EMRC, light purple).

